# sorAI - a suite of machine learning models to predict plant polyploidy from target capture data

**DOI:** 10.64898/2026.11.10.765735

**Authors:** Fahimeh Rahimi, Jessie A. Pelosi, J. Gordon Burleigh, Juannan Zhou

## Abstract

**Premise:** Polyploidy is a persistent feature of land plant evolution, yet it is often difficult to determine the ploidy of individual plants. The growing abundance of target capture data and other high throughput sequence datasets across plant species has generated interest in methods to infer ploidy from these data. We leverage chromosome count data and target capture data from the GoFlag probe set in ferns to train new machine learning models to infer if a sample is from a diploid or polyploid genome.

**Methods and Results:** We trained a convolutional neural network (CNN) for each GoFlag probe locus and developed ensemble models that use two loci and a random forest model that leverages all the loci to classify fern samples as diploid or polyploid. We tested the performance of these models using held-out test datasets of GoFlag target capture data across ferns and an independent test set from the genus *Lygodium*. We also performed experiments to evaluate the features of loci and sequences most associated with model performance and tested the performance of the models as sequence depth decreased.

**Conclusions:** These models, collectively named sorAI, outperform the frequency-based methods nQuire and nQuack on held-out test datasets of GoFlag target capture data across ferns. These models are capable of leveraging positional information and covariation among nucleotide sites beyond individual-site allele frequencies, and they perform well even with low sequence coverage. The random forest model, which uses all loci, appears to be robust to varying dataset features and achieves an accuracy of approximately 71–78% across test datasets and sequencing depths. sorAI is available at https://github.com/Fahimehgit/sorai, and the trained models are available at https://huggingface.co/Fahimehr140/sorai.

## INTRODUCTION

Polyploidy is a persistent feature of the evolution of land plants, and botanists have been long interested in the role of polyploidy in speciation, adaptation, and the generation of novel features in plants. Such studies require knowledge of the ploidy level of a diversity of plant samples. Understanding the ploidy also is critical for accurately genotyping plant samples. Yet, determining the ploidy of plant samples often is difficult and requires different lines of evidence.

One of the most direct ways to determine the ploidy level of a plant is by comparing the chromosome numbers among closely related taxa. There have been numerous efforts to assemble chromosome count data for plant species (e.g., Manton, 1950; Darlington and Wylie, 1956; Henniges et al., 2023; Rice and Mayrose, 2023; Simon et al., 2023). Although chromosome counts exist now for thousands of plant species, these still represent only a small fraction of total extant plant diversity. Furthermore, chromosome counting can be challenging in some plants, and usually requires freshly collected tissue, making it difficult to obtain from historical (i.e., herbarium) collections. Ploidy levels also can be inferred from genome size estimates, generally using flow cytometry (see Dolezel and Bartos, 2005; Kron et al., 2007) or measurements of cell size (e.g., guard cells in stomata, spores, or pollen; see Stebbins, 1950). However, these indirect correlates of ploidy can be highly variable among closely related taxa with the same ploidy and thus difficult to interpret or potentially misleading.

The availability of large genomic datasets offers a new source of data from which to infer ploidy levels. With high coverage genome sequence data, ploidy can be inferred without a reference sequence from *k*-mers, or strings of nucleotides of length *k* (e.g., Ranallo-Benavidez et al., 2020). While there is a rapidly growing number of nearly full plant genome sequences, they still are unavailable for the vast majority of plant species. Ploidy levels also can be inferred from the read frequencies of variable (often biallelic) sites. In these approaches, the high throughput sequencing (HTS) reads first are mapped to reference sequences. The ploidy of samples is then inferred by comparing the observed read frequencies to either expected ratios predicted for different ploidy levels or frequencies observed from simulations of sequences from different ploidy levels. For example, in a diploid each allele should be present in approximately 1/2 of the reads, and in a biallelic triploid site, one allele should be present in 2/3 of the reads, and the other in ⅓. The first software to use this approach was ploidyNGS, which enabled users to evaluate ploidy levels by visually inspecting allele frequency histograms (Augusto Corrêa dos Santos et al., 2017). Later, nQuire enabled users to infer ploidy using a likelihood approach based on the fit of read frequencies at biallelic sites to a Gaussian (i.e., Normal) mixture model (Weiß et al., 2018), and it has been incorporated in protocols designed explicitly to infer ploidy from plant phylogenomic datasets (Viruel et al., 2019, 2023). More recently, nQuack expands upon the methods from nQuire, using a more elaborate statistical framework for evaluating ploidy levels and including arguably more appropriate distributions for the mixture models (Gaynor et al., 2024). Tiley et al. (2023) also developed a variant of nQuire which uses different mixture models and is implemented as part of the PATÉ pipeline for phasing allelic or homeolog sequences (Tiley et al., 2024).

While the allele frequency approaches for inferring ploidy can be applied to many types of HTS data, in plants they most often are used with targeted capture data, and some protocols are explicitly designed for these data (e.g., Viruel et al., 2019, 2023; Tiley et al., 2023). Targeted capture is an approach to obtain data from hundreds of nuclear loci at a relatively low cost (e.g., Cronn et al., 2012; Weitemier et al., 2014; McKain et al., 2018). It generally uses short RNA probes, corresponding to selected loci, to bind to DNA from sequencing libraries. The bound DNA is then sequenced, generating a sequence dataset enriched for the chosen target loci. In plants, probe sets have been designed for extremely large groups of plants, like the Compositae (Mandel et al., 2014; Moore-Pollard, 2024), angiosperms (Johnson et al., 2019), flagellate land plants (Breinholt et al., 2021), the nitrogen-fixing clade within the rosids (Folk et al., 2021), and orchids (Eserman et al., 2021). Because targeted capture usually uses short-read sequencing, it can be effective on old or degraded DNA that may come from herbarium samples (Brewer et al., 2019; Forrest et al., 2019; Kates et al., 2021). Thus, targeted capture has been used to generate genomic sequences from a rapidly expanding proportion of plant species. While the allele frequency models appear to work well on targeted capture data in some cases (e.g., Gaynor et al., 2024), in others their accuracy has been questioned. In an extreme case, Pelosi et al. (2024) found that the accuracy of nQuire was <30% among samples of vining ferns (*Lygodiaceae*). This poor performance may be due to either the quality or quantity of the sequence data (e.g., inadequate read depth), and not necessarily inherent flaws in the methods; however, these results suggest that the allele frequency approaches may not perform well on real world plant datasets derived from targeted capture sequencing.

The availability of chromosome counts (e.g., Rice and Mayrose, 2023) and targeted capture datasets from thousands of taxa (e.g., Baker et al., 2022) suggests the possibility of a machine learning approach to estimate ploidy. In such a method, instead of comparing aspects of the sequence data (e.g., read counts for different alleles) to theoretical predictions, the machine learning model can be trained on targeted capture datasets of known ploidy and then deployed to predict the ploidy levels for novel samples. Such models would not be limited to bi-allelic frequencies or theoretical predictions, and they may be able to “learn” diagnostic features of ploidy level from the complex, real datasets, such as biases in the allele frequencies due to probe design. They also could incorporate features of the dataset not typically included in allele frequency analyses: for example, they could account for neighboring sites instead of treating each nucleotide site independently, thereby potentially leveraging the distribution of (micro-)haplotypes for prediction. In this study, we describe a new approach that uses machine learning to estimate the ploidy of a sample from raw sequencing reads generated from targeted capture, and we test it with a dataset of ferns. Ferns are a clade of more than 10,000 species (PPG I, 2016) that have long been a model system for studying polyploidy in plants (Manton, 1950; Klekowskiand Baker, 1966); an estimated 30% of fern speciation events are associated with a change in ploidy (Wood et al., 2009). The GoFlag 408 flagellate land plant probe set (Breinholt et al., 2021) has been used to generate data from thousands of fern samples, which have been used in numerous evolutionary studies (e.g., Fawcett et al., 2021; Lima et al., 2023; Mendez Reneau et al., 2023, 2024; Koubinová et al., 2024; Pelosi et al., 2024; Sigel et al., 2025). We assembled a set of targeted capture data from fern samples from species with ploidy estimates from chromosome counts. We used these data to train new machine learning models to infer if a fern sample is diploid or polyploid. We evaluate the performance of these new models and compare their performance to that of existing methods to infer ploidy. Overall, we found that our models outperform the frequency-based nQuire and nQuack on held-out test datasets, and that the random forest model, which aggregates predictions across all loci, also outperforms both methods on an independent test set consisting of samples from the genus *Lygodium*. We further performed model interpretation tasks and showed that our models are capable of leveraging positional information and covariation among nucleotide sites beyond individual-site allele frequencies.

## METHODS

### Sampling and Data

First, we obtained a list of chromosome counts from fern species from the Chromosome Counts Database (CCDB; Rice and Mayrose, 2023) and used the chromosome counts to determine which species were diploid or polyploid by comparing chromosome numbers among closely related taxa. We did not distinguish among different types of polyploids (e.g., tetraploids, hexaploids, or octoploids), nor did we distinguish whether the polyploids were allopolyploids or autopolyploids. We then took a list of the species that we determined to be unambiguously diploid or polyploid based on chromosome counts (i.e., species in which all available chromosome counts were consistent with the ploidy designation), and identified samples that the GoFlag project had sequenced using the GoFlag 408 or 451 flagellate land plant probe set. The GoFlag 408 probe set is an optimized subset of the GoFlag 451 probe set (Breinholt et al., 2021), and for all the samples and analyses, we only considered loci from the GoFlag 408 probe set. Each locus in these probe sets represents an exon on a single or low copy nuclear gene that is at least 120 base pairs (bp) in length. In total, we found targeted capture sequence datasets from 640 samples generated using the GoFlag 408 flagellate land plant probe set and six samples generated using the GoFlag 451 probe set. Among the chosen samples, the targeted capture sequencing recovered an average of 371 loci, and the reference sequences for the 384 loci used in our analyses have an average length of 190 bp (range: 111–1,068 bp).

### Overall workflow

The overall workflow, illustrated in Figure 1A, begins with quality control and preprocessing of raw target-capture reads. Processed samples were then randomly partitioned into non-overlapping training, model-selection validation, and held-out evaluation sets using an 80/10/10 split (N = 516, 64, and 66 samples, respectively; Figure 1B).

**Figure 1.**
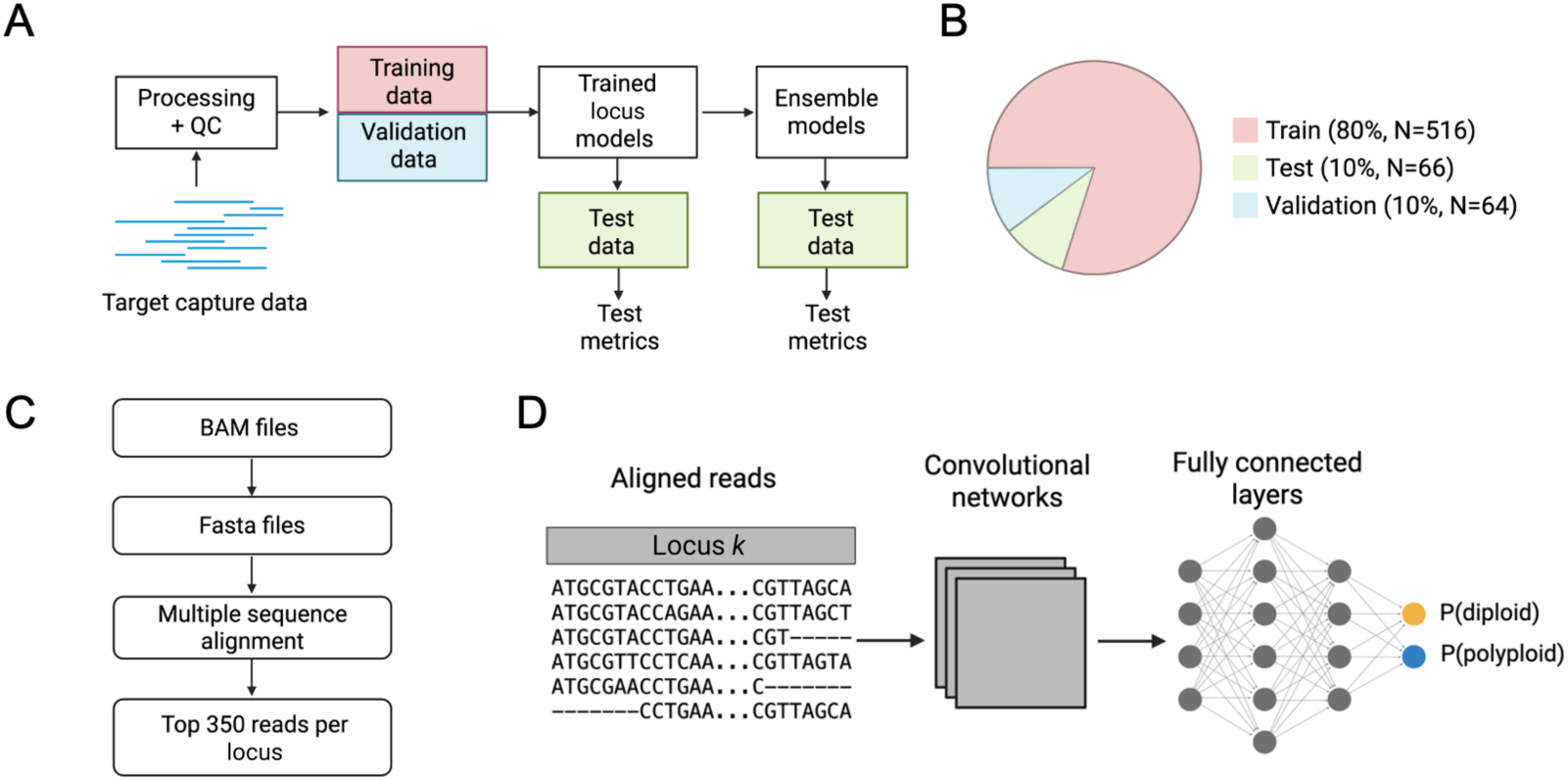
Machine learning models for identifying polyploids using target capture data. (A) Overall workflow of our method. The raw target capture reads acquired using the GoFlag flagellate land plant probe set are first processed and filtered. A subset of samples was used as training and validation data to train CNN models for individual loci. The trained CNN models are then aggregated for different pairs of loci to generate a series of ensemble models to further improve ploidy prediction accuracy. (B) A total of 646 samples were randomly split into training, validation, and test datasets. The training dataset was used to learn the weights of the neural network models. The validation dataset was used to select the hyperparameters specifying the CNN model architecture. The test dataset (N = 66) was further divided into a calibration subset (N = 32) and a final held-out test subset (N = 34; see Methods), and was never used during model training. (C) Bioinformatic pipeline for processing raw target capture reads. For each sample and each locus, only the 350 reads with the fewest gaps after alignment were retained. (D) Ploidy prediction model architecture. The input to the model consists of aligned target capture reads for a given locus. The input is then passed through several convolutional layers, and the output is passed through several fully connected layers to predict the probability that the sample is diploid or polyploid.

We first trained convolutional network (CNN) models, each based on the training data for a single locus. To improve predictive performance, pairs of locus-specific CNN models were later combined into ensemble models, retrained using the same training/validation framework and evaluated on the same independent test set, which was not used during model selection. We also similarly developed and tested a random forest model that uses all of the loci.

### Sequence alignment and preprocessing

For each sample we generated binary alignment map (BAM) files that map the sequence reads from the GoFlag targeted capture sequence datasets to reference sequences for each locus. First, we trimmed the FASTQ files with paired end raw sequence reads using Trim Galore (https://github.com/FelixKrueger/TrimGalore), which removes adapters and low-quality sequences. We then assembled sequences for each locus targeted by the GoFlag probe set using the pipeline described in Breinholt et al. (2021). In cases in which the GoFlag pipeline produced more than one sequence per locus, we chose as the reference the longest sequence, or the first sequence output if they are of equal length. Next, we mapped the trimmed paired end reads to the reference sequences for each locus using the Burrows-Wheeler Aligner (BWA; Li and Durbin, 2009), which outputs the alignment in a sequence alignment map (SAM) file. We then used SAMtools v1.3.1 (Li et al., 2009) to convert the SAM files into compressed BAM files. These are the same initial steps used to prepare data for analyses with nQuire (Viruel et al., 2023).

BAM files were processed into a standardized alignment format for CNN training (Figure 1C; Supplemental Figure 1). For each sample and locus, reads were converted from BAM to FASTA format. We used pysam (v0.9.1) for BAM processing, with SAMtools v1.3.1 (Li et al., 2009) as a fallback when needed. Read alignments were generated with MAFFT v7.520 (Katoh and Standley, 2013) unless alignment was explicitly skipped in accelerated runs. To standardize alignment structure, we removed internal gap columns — alignment columns corresponding to gap positions in the reference sequence, excluding its first and last positions — from all sequences, and separately trimmed all sequences to the window defined by the leading and trailing gap positions of the reference sequence. Both steps were keyed to the reference sequence rather than a percentage-based gap threshold across all reads. The retained gap characters therefore represent alignment gaps (indel-related positions) in the processed multiple-sequence alignments.

To construct fixed-size model inputs, we retained up to 350 reads per locus for each sample. When more than 350 reads were available, we selected the 350 reads with the fewest gap characters and ordered them by ascending gap count (fewest gaps first); when 350 or fewer reads were available, all reads were retained in their original order from the assembled FASTA file. When fewer than 350 reads were available, the remaining rows were padded with entire rows of the gap token so that each locus had the same read-depth dimension (350 reads plus the reference). Nucleotides were then numerically encoded using a fixed tokenization scheme (A = 1, C = 2, G = 3, T = 4, gap/other = 0). Each sequence — reference and reads alike — was standardized to a fixed length of 154 nucleotide positions, near the mean length of the recovered locus sequences (144.6 bp), by truncating sequences longer than 154 positions (retaining the first 154 positions) and right-padding shorter sequences with the gap token. This produced final tensors of 351 sequences (1 reference + 350 reads) × 154 positions per locus per sample.

### CNN model architecture and training

Samples were partitioned into four non-overlapping subsets: training (N = 516), model-selection validation (N = 64), held-out calibration (N = 32), and final test (N = 34). The training set was used to fit model parameters. The model-selection validation set was used during development for hyperparameter optimization. After hyperparameters were fixed, the held-out calibration set—kept fully separate from model fitting and hyperparameter tuning—was used for post-selection checks (e.g., threshold and probability-calibration behavior) before final reporting. The final test set was reserved for independent performance evaluation and was not used for model fitting, hyperparameter selection, or threshold tuning. In analyses requiring a larger held-out cohort, we additionally report results on the combined held-out set (held-out calibration + final test; N = 66).

For each locus, we trained a separate convolutional neural network (CNN) to classify samples as diploid or polyploid using aligned, tokenized read matrices (Figure 1D; Supplemental Figure 2). We did not attempt to distinguish among different polyploidy levels (e.g., triploid, tetraploid, hexaploid) due to the limited size of our training set. Locus-specific CNNs followed a LeNet-style architecture with 1–3 convolutional layers (16–32 filters; 3×3 or 4×4 kernels), ReLU/LeakyReLU/ELU activations, max or average pooling, and two fully connected layers (64–128 units) with shared dropout (0.1–0.3). A sigmoid output layer produced the predicted probability of polyploidy.

Model parameters were trained using Adam optimizer (Kingmaand Ba, 2014) under binary cross-entropy loss. Hyperparameters were selected independently for each locus using Optuna (Akiba et al., 2019; 25 trials per locus). The full search space and selected values are reported in Supplemental Table S1. The search space included architectural choices (number of convolutional layers, filter configuration, kernel size, activation, pooling, and fully connected width) and optimization parameters (learning rate: 10⁻⁵ to 10⁻³; weight decay: 10⁻⁵ to 10⁻³; epoch budget: 10–30). To improve robustness to read-order artifacts, the 350 non-reference reads were randomly permuted at each training epoch while the reference row remained fixed. Final performance was reported using accuracy and area under the receiver operating characteristic curve (AUROC) on the independent final test set, with combined held-out analyses (N = 66) reported where specified.

### Training models on pairs of loci (ensemble models)

In addition to single-locus CNN models, we developed ensemble models that combine information from pairs of loci to improve ploidy prediction (Figure 1A). We retained 384 loci as candidates for downstream ensemble modeling after preprocessing and model-quality filtering. Considering all pairwise combinations of these 384 loci would have required training and evaluating 384 × 383 / 2 = 73,536 pair models. Thus, to keep this analysis computationally tractable, we trained pair models for a subset of locus pairs and analyzed a random subsample of 1,996 of these pair models. This subsample included every one of the 384 candidate loci at least once (1–27 pairs per locus; mean = 10.4, median = 9), and per-locus representation was independent of single-locus validation AUROC (Pearson r = 0.04, Spearman ρ = 0.05), indicating that the sampled pairs were not biased toward higher-or lower-performing loci and instead spanned the full range of individual locus quality (validation AUROC range: 0.573–0.960, mean = 0.811). This subsample allowed us to evaluate ensemble performance both for pairs of individually strong loci and — as illustrated below — for pairs where one or both constituent loci performed only moderately or near chance on their own.

Rather than combining the raw sequencing reads from two loci directly, each ensemble model combines the internal, learned summary that each single-locus CNN model produces for a sample — a compact numerical representation of the patterns that model uses to distinguish diploids from polyploids — and uses this combined summary as input to a second-stage classifier trained specifically on that pair of loci. Specifically, for each candidate locus, we extracted the final hidden CNN representation (i.e., output by the layer immediately before the sigmoid activation) and trained a locus-specific projection head to map this representation to a common 500-dimensional embedding. This was necessary because locus-specific CNN configurations can produce representations of variable sizes due to different CNN model hyperparameters. Each projection head consisted of an embedding block (a fully connected layer to 500 dimensions, followed by ReLU activation and dropout [p = 0.2]) and a classifier block (fully connected layers of 256 and 1 units, with ReLU activation and dropout [p = 0.2] on the first layer). The models were trained with a binary cross-entropy loss (weighted by class imbalance). Projection heads were trained independently for each of the 384 candidate loci, using AdamW (learning rate = 1×10⁻³, weight decay = 1×10⁻⁴) for up to 200 epochs with early stopping (patience = 20 epochs) and a batch size of 64.

For each pair of loci, we discarded the classifier block of each projection head and used only its embedding block, so that each locus in the pair was represented by a 500-dimensional embedding for each sample. The two embeddings were then concatenated into a single 1,000-dimensional feature vector, which served as the input to the pair-level classifiers. We evaluated four classifier heads on these pair features, each with a fixed configuration rather than pair-specific hyperparameter tuning: logistic regression (L-BFGS solver, max_iter = 2000, with feature scaling); support vector machine with a radial basis function kernel (C = 10, gamma = "scale", with feature scaling); gradient boosting (300 estimators, learning rate = 0.05, max depth = 3); and a multilayer perceptron (two hidden layers of 256 and 128 units, alpha = 1×10⁻³, initial learning rate = 1×10⁻³, max_iter = 2000, with early stopping and feature scaling). For each pair, all four heads were fit on the training split, and the best-performing head was selected by AUROC on the model-selection validation split; that head was then refit on the combined train+validation data and evaluated on the held-out calibration and final test sets. This procedure yielded 1,996 locus-pair model datapoints, with the best-performing head selected as logistic regression (n = 239), SVM-RBF (n = 252), gradient boosting (n = 698), or MLP (n = 807) in different pairs.

All analyses were run on Linux with Python 3.9, using PyTorch for model training/inference, Optuna for hyperparameter optimization, and scikit-learn for ensemble classifiers. Runs used fixed random seeds and fixed data splits for reproducibility, with GPU when available and CPU fallback otherwise. Training the reported pairwise models required approximately 12.8 GPU-hours in total (mean 23.1 s per pair; range 3.5–72.2 s per pair).

### Random Forest (RF) model using all loci

In addition to the CNN and ensemble models, we also developed an ad hoc Random Forest (RF) model that aggregates signals across all available loci for each sample. This may be useful in cases when single-locus CNN predictions are poorly calibrated for the samples or have low individual accuracy. We trained the RF model using the original fern training data. Specifically, each sample was represented using all available locus-level CNN probabilities for that sample (i.e., no fixed locus panel was required per sample). From this per-sample probability profile, we computed summary descriptors of signal strength and uncertainty, including central tendency (average behavior), dispersion (spread across loci), confidence structure (how strongly predictions deviate from 0.5), and entropy (overall prediction uncertainty). The RF model uses these descriptors to produce a final ploidy probability, allowing each sample to contribute evidence from all usable loci despite uneven locus recovery.

### Evaluation and Reproducibility

For final classification, probabilities were converted to binary labels using predefined thresholds (0.5 for CNN/ensemble/hybrid; fixed RF threshold from training-only calibration). We compared this probability-averaging framework against alternative decision rules (including majority voting) and retained probability averaging for CNN/ensemble/hybrid because it yielded the best held-out performance in our comparisons. Performance of all CNN and ensemble models was evaluated on the independent test set (19 diploids and 15 polyploids) using accuracy, defined as the proportion of samples correctly classified, and the AUROC, which measures the ability to distinguish diploids from polyploids across all possible classification thresholds. AUROC complements accuracy by evaluating overall discriminatory ability independently of a single classification threshold and is generally less sensitive to class imbalance.

### Comparison with Existing Ploidy-Inference Methods

To benchmark performance, we compared our approach with nQuirev20180405 (Weiß et al., 2018) and nQuack v1.0.0 (Gaynor et al., 2024), which we applied to the same sequence data from the 646 samples. For nQuire, analyses were run with and without de-noising, and ploidy was inferred from the model with the lowest Δlog-likelihood relative to the free model. Diagnostic metrics such as sum of squared residuals (SSR), regression slope, and R² were used to assess fit quality. For nQuack, we followed the published tutorial, evaluating 18 model configurations spanning six probability distributions and three mixture model types. Both nQuire and nQuack estimate the ploidy level of polyploids, but for our comparisons we only considered if the result was diploid or polyploid.

We also phased the reads from all of the samples using PATÉ (Tileyet al., 2024) to determine if the estimated proportion of variable loci or overall heterozygosity of each sample could be used to distinguish between diploids and polyploids. PATÉ requires an input file that has the ploidy for all samples and limits the number of possible alleles. We ran PATÉ on all samples assuming that they were all diploid, which is a reasonable strategy if the ploidy levels were unknown. We used the same reference sequences from each locus that we used to generate the original BAM files.

### Exploratory analyses of trained CNN models

A key distinction between our CNN framework and allele-frequency methods (e.g., nQuire and nQuack) is that CNNs can leverage inter-site covariation via convolutional filters and nonlinear layers. To test whether our models have learned to exploit this information, we applied three shuffling perturbations to reads from a held-out evaluation set of 66 samples (32 calibration samples plus 34 test samples): (1) global row shuffling, in which rows in the read alignments were permuted consistently across columns, preserving per-read composition while perturbing row-wise structure relative to the reference; (2) column shuffling, in which alignment columns were permuted as intact units, disrupting positional order while preserving within-column nucleotide composition; and (3) independent within-column row shuffling, in which the nucleotides were shuffled independently within each column, preserving per-column allele frequencies while disrupting covariation across positions (Figure 4A). We applied these perturbations to test data for the top 100 individual single-locus CNN models and the top 100 ensemble pair models. For each model and each perturbation, shuffled-input performance was compared with unshuffled-input performance on the same held-out samples. Performance declines under these perturbations provide evidence that predictions depend on aspects of alignment structure beyond the marginal per-site nucleotide or allele-frequency distributions preserved by all three shuffling procedures.

### Correlational study between model performance and data features

To evaluate how properties of the input-alignment relate to model performance, we tested associations between per-locus test AUROC and summary statistics computed from the multiple-sequence alignments used to train each of the 384 locus-specific CNN models. For each locus, we computed three metrics: (1) gap fraction, the proportion of gap characters across all alignment cells (reads × positions); (2) variant density, the proportion of alignment columns with more than one observed nucleotide state after excluding gaps; and (3) mean Shannon entropy, the average per-column entropy across alignment positions, where larger values indicate greater within-column nucleotide diversity averaged across positions. We then quantified associations between each metric and test AUROC across loci using both Pearson and Spearman rank correlation.

### Effect of read depth on accuracy of CNN and RF models

To assess the sensitivity of our models to sequencing depth and determine how reduced read coverage affects predictive performance, we evaluated both single-locus CNN and RF models using the same depth-reduction procedure. Specifically, for each sample and depth fraction (1, 1/2, 1/4, 1/8, 1/16, and 1/32 of full depth), the reference sequence was retained, non-reference reads were randomly subsampled without replacement to the target depth, and the retained non-reference reads were randomly permuted to minimize order-dependent bias; four replicate resamples were generated per depth fraction using fixed random seeds.

For the CNN model analysis, we applied this procedure to the 10 top-performing CNN locus models, evaluating each fixed, pretrained model directly on the depth-reduced reads from the held-out N=34 test set without retraining. For the RF analysis, we applied the same depth-reduction procedure across all available loci, recomputed each locus’ CNN probability from the depth-reduced reads, transformed these into the RF model’s feature set, and passed them through the fixed, pretrained RF bundle. In both cases, performance at each depth fraction was summarized as the mean ± SD across the four replicates for test accuracy and AUROC.

### Application to an independent dataset (*Lygodium)*

To evaluate the models on fern samples not represented in model training, we applied our trained model with frozen weights to a published dataset of target-capture samples from 35 species of *Lygodium* (Lygodiaceae), generated with the GoFlag 408 probe set (Pelosi et al., 2024). The GoFlag probes recovered an average of only 266 loci per sample, compared to an average of 371 loci for all training set samples, therefore representing a challenging real-world dataset. This dataset included 60 samples from 18 species that had ploidy annotations based on chromosome counts and were not included in the original training set used to develop the ML models. Among these samples, 41 were diploid and 19 were tetraploid. Within the tetraploid subset, 10 samples were classified as likely autotetraploid (*L. flexuosum*, *L. japonicum*, and *L. longifolium*) and 6 as allotetraploid (*L. cubense*, *L. heterodoxum*, and *L. oligostachyum*). The remaining samples from this study had ambiguous (e.g., multiple cytotypes) or missing ploidy annotations and were not included in supervised accuracy/AUROC analyses.

We used four strategies to predict ploidy in the *Lygodium* samples. First, we applied the ten highest-ranked single-locus CNN models to each sample. Second, we applied five top-ranked locus-pair ensemble models (loci 146+389, 146+320, 139+146, 16+342, and 158+373), each with its selected classification head (SVM-RBF for four pairs and MLP for 139+146). Third, we evaluated a hybrid strategy that combines the single-locus CNN models with a locus-pair ensemble model. For a given sample, the hybrid probability of polyploidy was calculated as the mean of two components: the predicted probabilities from the 10 single-locus CNN models and the predicted probability from one locus-pair ensemble model. Because this second component depends on which ensemble pair is chosen, the hybrid probability is not a single fixed value — it changes depending on which locus pair is included. To illustrate this, we computed the hybrid probability separately using each of the five top-ranked locus-pair ensemble models described above, producing five hybrid predictions per sample rather than one. This allowed us to directly compare how the choice of ensemble pair influences the combined CNN + ensemble prediction. Finally, we also used the ad hoc Random Forest (RF) model that aggregates signals across all available loci for each sample.

To examine whether model performance differs between autopolyploid and allopolyploid samples, we subset the *Lygodium* validation samples into two comparisons — autotetraploid vs. diploid and allotetraploid vs. diploid — using the classifications of Pelosi et al. (2024). For each comparison and each model type (CNN, ensemble, hybrid, and RF), we performed 5,000 non-stratified bootstrap resamples, sampling individual samples with replacement from the combined diploid and polyploid-subtype subset. For each resample, we computed accuracy (at the 0.5 classification threshold) and AUROC; bootstrap draws containing samples from only one class were discarded from the AUROC calculation, as AUROC is undefined in that case. Reported accuracy and AUROC values are the mean of the bootstrap distribution, and 95% confidence intervals were calculated as the 2.5th and 97.5th percentiles of that distribution.

## RESULTS

### Model Performance of CNN individual locus-based models

We evaluated all 384 locus-specific CNN models on the held-out test set (N = 34; Figure 2, Supplemental Table S1). Across models, the mean accuracy was 0.641 ± 0.067 (mean ± SD; Figure 2A), and the mean AUROC was 0.729 ± 0.071 (Figure 2B). The model based on locus 238 achieved the highest AUROC (0.923; accuracy = 0.824; Figure 2C).

**Figure 2.**
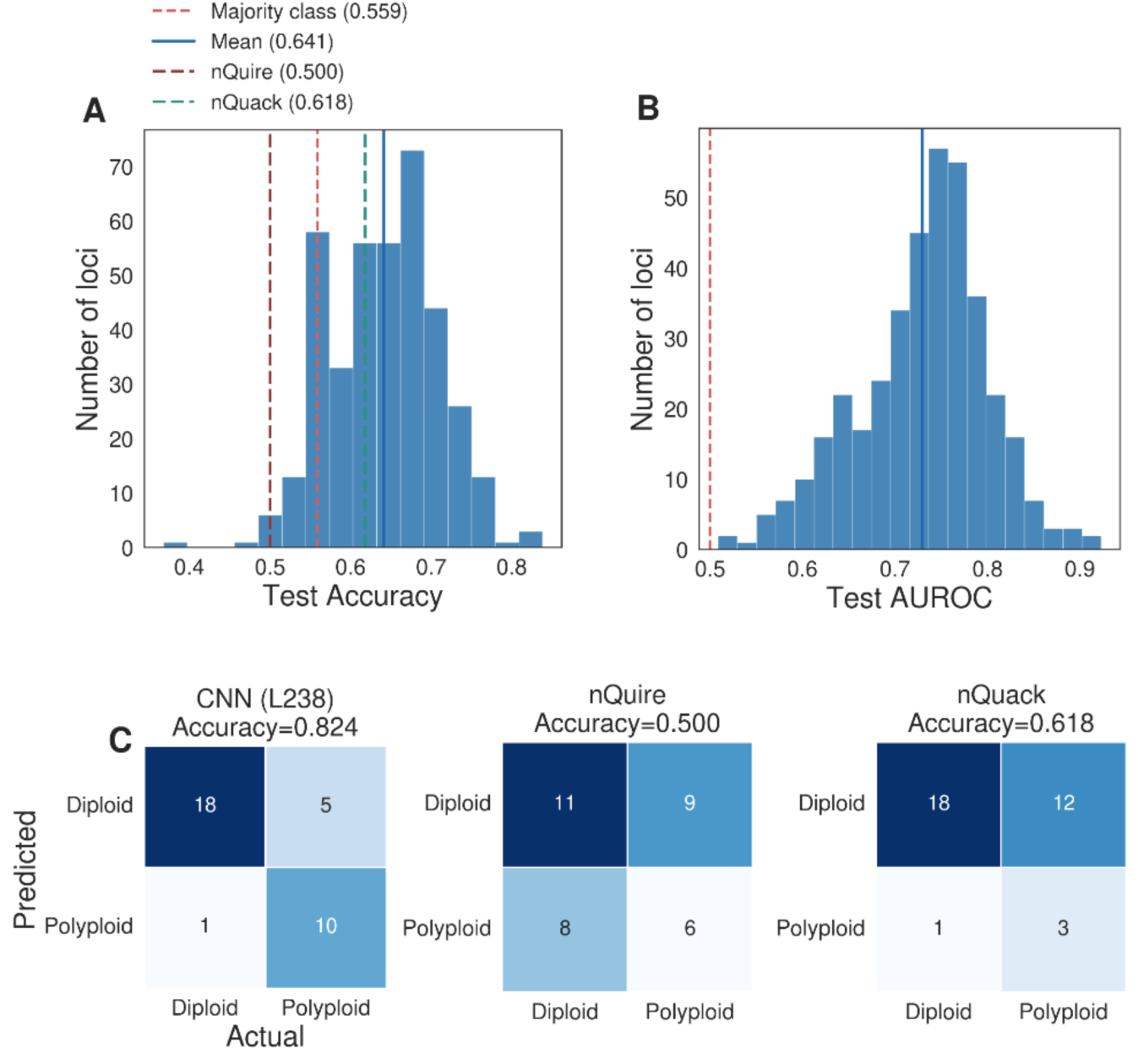
Distribution of performance metrics across all 384 individual CNN locus models evaluated on an independent test set of 34 samples (19 diploid and 15 polyploid). (A) Histogram of test accuracy across all single-locus CNN models; each bar corresponds to one possible accuracy value on the 34-sample test set (k/34). The red dashed line indicates the majority-class baseline (0.559, the proportion of diploid samples), and the dark red and teal dashed lines indicate the accuracy of nQuire (0.500) and nQuack (0.618), respectively. (B) Histogram of test AUROC across all single-locus CNN models; the red dashed line indicates random classification performance (0.50). In (A) and (B), solid blue lines indicate the mean across loci. (C) Confusion matrices for the best-performing single-locus CNN model (L238), nQuire, and nQuack on the 34-sample test set. Across all loci, the mean test AUROC was 0.729 ± 0.071 and the mean test accuracy was 0.641 ± 0.067 (mean ± SD). The best-performing model (L238) achieved a test accuracy of 0.824 and an AUROC of 0.923.

### Ensemble Model Performance

To determine whether integrating information across loci improves ploidy prediction, we trained and evaluated ensemble models on pairs of loci using four classification heads (logistic regression, SVM-RBF, gradient boosting, and MLP; see Methods). In total, we evaluated 1,996 locus-pair model configurations. Among the 1,996 analyzed pairs, the best-performing model with respect to AUROC combined loci 146 and 313 with an MLP head and achieved an AUROC of 0.993 and an accuracy of 0.912 on the held-out 34-sample test set. To illustrate the magnitude of the ensemble effect even when constituent loci are individually weak, Figure 3A and B compares a representative ensemble model (loci 15 and 350, gradient boosting head; accuracy = 0.941, AUROC = 0.963) to its two constituentsingle-locus CNN models (locus 15: accuracy = 0.706, AUROC = 0.793; locus 350: accuracy = 0.588, AUROC = 0.740) and to the best single-locus CNN model overall (locus 238: accuracy = 0.824, AUROC = 0.923). Although neither locus 15 nor locus 350 performs well individually, and locus 350 in particular has an accuracy only modestly above random chance, their combination substantially outperformed both constituents and even the best individual CNN locus. To assess this pattern across all 1,996 evaluated configurations, we compared the performance of each locus-pair ensemble to the stronger of its two constituent single-locus models (Figure 3C, D). Across all four classifier heads (logistic regression, n = 239; SVM-RBF, n = 252; gradient boosting, n = 698; MLP, n = 807), a large majority of locus-pair models fell above the y = x reference line for both accuracy and AUROC, indicating that most ensembles outperformed the best of their two constituent loci. Across all 1,996 locus pairs, 92.33% of points in the accuracy panel and 93.74% of points in the AUROC panel lie above the identity line. This pattern was consistent across classifier types, suggesting that integrating information across loci captures complementary discriminatory signals not available from either locus alone, regardless of which classification head is used to combine them.

**Figure 3.**
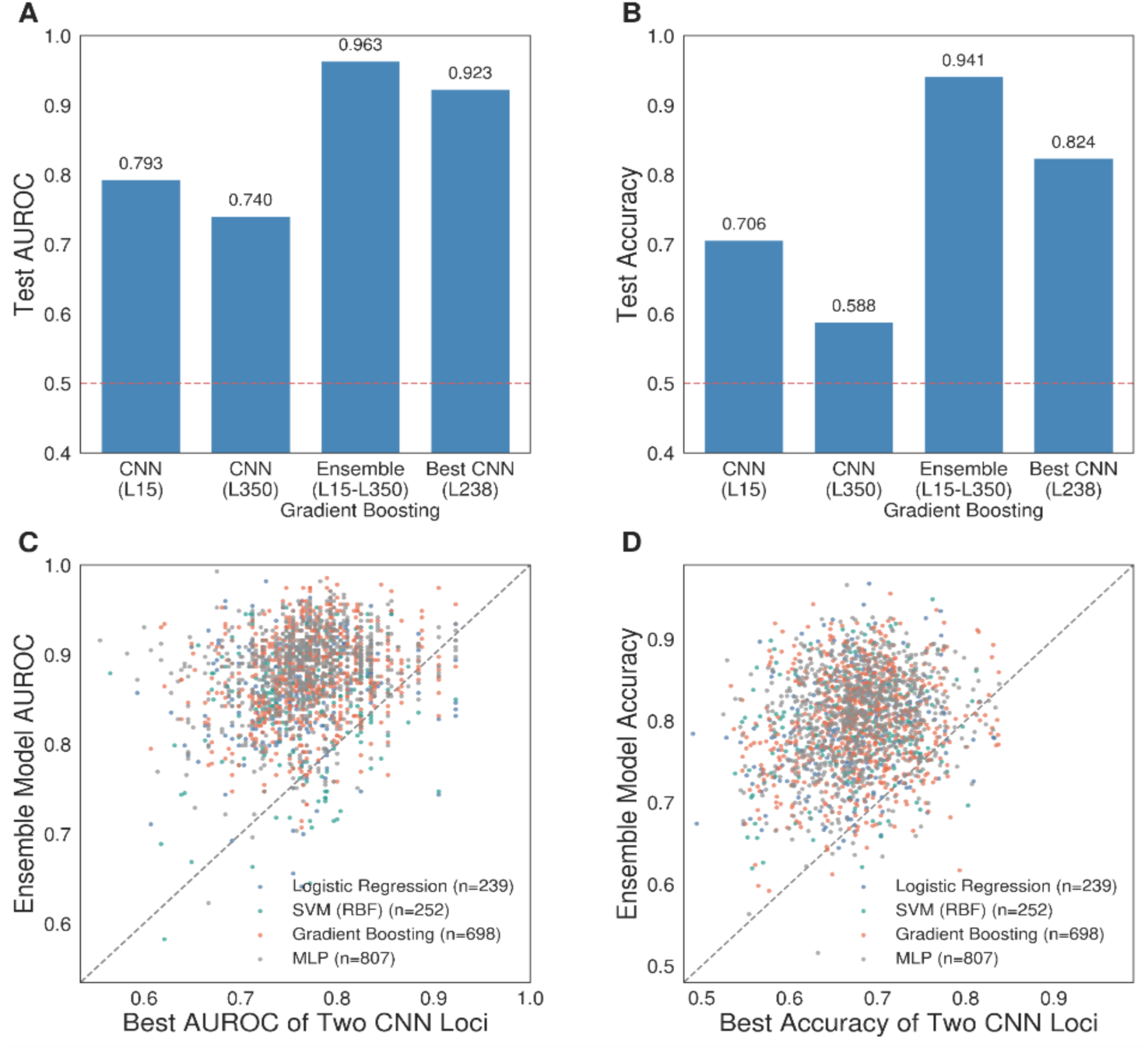
Ensemble models outperform individual single-locus CNN models on the independent test set. (A, B) Comparison of test AUROC (A) and test accuracy (B) for a representative ensemble model (loci L15 and L350, gradient boosting head) against its two constituent single-locus CNN models and the best-performing single-locus CNN model overall (L238). This pair illustrates that a moderately performing locus (L15: AUROC = 0.793, accuracy = 0.706) combined with a weaker locus (L350: AUROC = 0.740, accuracy = 0.588) can yield an ensemble (AUROC = 0.963, accuracy = 0.941) that substantially outperforms both constituents and exceeds even the best individual locus model (L238: AUROC = 0.923, accuracy = 0.824). Red dashed lines indicate random classification performance (0.50). (C, D) Scatter plots comparing the AUROC (C) and accuracy (D) of each of 1,996 locus-pair ensemble models (y-axis) to the better-performing of its two constituent single-locus CNN models (x-axis), across four classification heads: logistic regression (n = 239), SVM-RBF (n = 252), gradient boosting (n = 698), and MLP (n = 807). Points above the diagonal indicate ensembles that outperform the stronger of their two constituent loci. Across all 1,996 locus pairs, 92.33% of points in the accuracy panel (D) and 93.74% of points in the AUROC panel (C) lie above the identity line, indicating that ensemble performance exceeds the single-locus baseline for the large majority of pairs.

### Random Forest Model Performance

For comparison, we also evaluated the RF model on the original 34-sample held-out test set, where it achieved an accuracy of 0.765 (26 of 34 samples correctly classified) and an AUROC of 0.870. This is higher than the average single-locus CNN model (accuracy = 0.641, AUROC = 0.729) but lower than the best-performing single-locus CNN model (accuracy = 0.824, AUROC = 0.923).

### Comparison with Other Methods

To benchmark our approach against existing ploidy-inference methods, we applied nQuire (Weiß et al., 2018) and nQuack (Gaynor et al., 2024) to the same 34 independent test samples used above. For nQuire, we used the ‘lrdmodel’ output and assigned each sample to the ploidy class—diploid, triploid, or tetraploid—with the lowest Δlog-likelihood relative to the free model, following the authors’ recommendations. We performed this analysis using both full and denoised BAM files. Both input types yielded an accuracy of 50.0% (17 of 34 samples correctly classified as diploid or polyploid; Figure 2A, C). We also examined nQuire fit diagnostics (sum of squared residuals [SSR], regression slope, standard error, and R²). Only six samples received the same ploidy call across all four diagnostics. SSR and slope showed the lowest overall accuracy (both 41.18%) but with opposite biases: SSR classified most samples as polyploid (sensitivity, the proportion of polyploids correctly identified: 80.00%; specificity, the proportion of diploids correctly identified: 10.53%), whereas slope favored diploid calls (sensitivity 26.67%, specificity 52.63%). Standard error and R² performed somewhat better (both 58.82%), but standard error classified nearly all samples as diploid (sensitivity 6.67%, specificity 100%), whereas R² was more balanced (sensitivity 53.33%, specificity 63.16%).

For nQuack, we evaluated the 18 model combinations recommended in the published workflow. For each sample and configuration, the predicted ploidy was taken as the class with the lowest ΔBIC. Correct classifications ranged from 15 to 21 of 34 samples (44.12%–61.76%). The best-performing configuration (normal-uniform, fixed_2, where only variance is free) achieved 61.76% accuracy (Figure 2A), outperforming nQuire on this test set. However, this model showed a strong diploid bias: it correctly classified 18 of 19 diploid samples (94.74% specificity) but only 3 of 15 polyploid samples (20.0% sensitivity), indicating that its overall accuracy advantage over nQuire reflects better performance on diploids rather than reliable polyploidy detection.

Additionally, phasing-based summaries from PATÉ indicated that, on average, polyploid samples had higher heterozygosity, a higher proportion of variable loci, and a higher proportion of phased loci than diploid samples (Supplemental Figure 7). However, these distributions overlapped substantially (Supplemental Figure 7), indicating that these summary statistics alone are not sufficient as standalone ploidy diagnostics.

### Single-locus CNN models rely on positional structure, while ensemble models additionally rely on inter-site covariation

We applied the three shuffling perturbations (see ‘Exploratory analyses of trained CNN models’ in Methods) to the top 100 individual CNN locus models: global row shuffling (disrupting row-wise structure), column shuffling (disrupting positional order), and independent within-column row shuffling (disrupting inter-site covariation while preserving per-column allele frequencies; Figure 4A).

**Figure 4.**
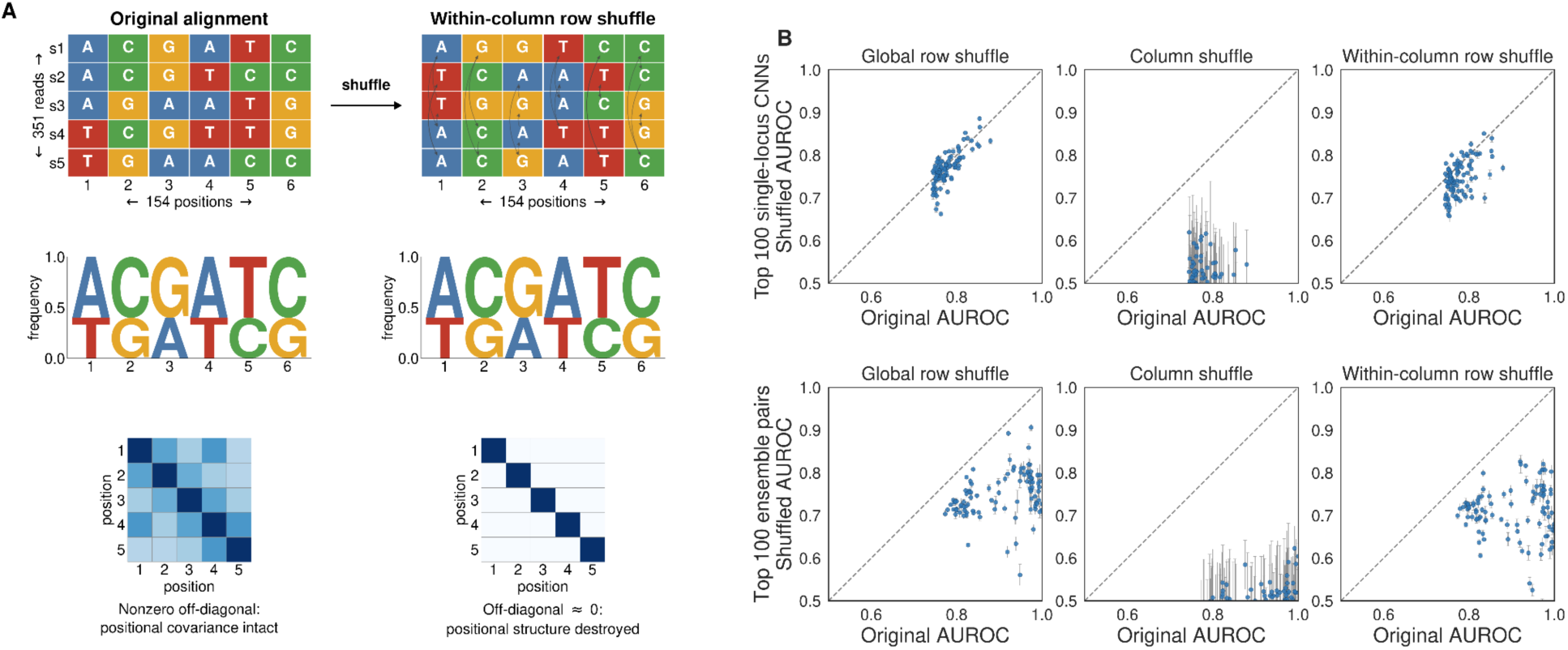
Models rely on positional structure, and ensemble models additionally capture inter-site covariation. (A) Schematic of the independent within-column row shuffling procedure: nucleotides within each alignment column are shuffled independently across reads, preserving per-position (per-column) allele frequencies while destroying covariation between positions. **(B)** Comparison of model performance (AUROC) before and after each of three shuffling perturbations — global row shuffling (left), column shuffling (middle), and independent within-column row shuffling (right, corresponding to the schematic in A) — applied to held-out reads from a 66-sample evaluation set (32 calibration + 34 test samples). Top row: the top 100 individual single-locus CNN models. Bottom row: the top 100 ensemble pair models. Points falling below the diagonal indicate reduced performance after shuffling relative to the unshuffled control. Across both single-locus and ensemble models, column shuffling produced the largest performance decline, indicating strong dependence on positional order; ensemble models additionally showed substantial degradation under within-column row shuffling, indicating reliance on inter-site covariation not evident in single-locus models.

Performance of the CNN models was largely unaffected by row shuffling (mean accuracy change: −0.004; mean AUROC change: +0.008), but substantially degraded by column shuffling (mean accuracy drop: 0.150; mean AUROC drop: 0.270) (Figure 4B, top row), indicating that the single-locus models depend heavily on the positional identity of alignment columns while remaining insensitive to the order in which reads are presented, consistent with the random per-epoch read permutation used during training (see Methods). Notably, we also observed a moderate drop in model performance (mean accuracy drop: 0.014; mean AUROC drop: 0.031) when rows were shuffled independently within each column, which disrupts the covariation between positions. A subset of locus models wereinsensitive to this procedure, but others suffered a performance drop of up to 10% in AUROC — evidence that at least some of the CNN models have learned to exploit covariation between positions when making predictions.

We then applied the same procedures to the top 100 ensemble pair models. Unlike the single-locus models, the ensemble models showed a drop in performance under row shuffling (mean AUROC dropped by 0.157), indicating that these models rely on the ordering of sequencing reads for prediction. This behavior is expected as the ensemble models were trained on the concatenated read embeddings of the two loci calculated based on fixed read ordering, instead of shuffled reads like the CNN models. Therefore, future applications should retain the existing procedure of ordering reads by ascending gap count (fewest gaps first) to ensure high prediction performance. Column shuffling produced an even larger drop in performance than for the single-locus models (mean AUROC dropped by 0.402). Finally, the performance drop under independent row shuffling was more pronounced for the ensemble models (AUROC drop = 0.197), suggesting that these models rely more meaningfully on covariation between nucleotide positions in addition to positional structure.

### Characteristics of loci influencing CNN prediction performance

To examine how sequence characteristics influence CNN model behavior, we quantified associations between locus-level performance and alignment-derived features across all 384 loci (Figure 5). Across the 384 loci, gap fraction showed the strongest relationship with AUROC (Figure 5A; Pearson r = −0.41, p < 1×10⁻⁴; Spearman ρ = −0.36, p < 1×10⁻⁴), indicating that alignment gaps substantially reduce discriminatory performance. Variant density showed inconsistent evidence of association with AUROC (Figure 5B): the Pearson correlation was weak but statistically significant (r = 0.17, p = 0.00113), while the corresponding Spearman rank correlation was weak, negative, and not statistically significant (ρ = −0.06, p = 0.238), indicating no robust monotonic relationship between variant density and AUROC. Mean per-position Shannon entropy — a measure of how evenly nucleotides are distributed across the four possible bases at a given alignment column, with higher values indicating greater local sequence diversity among the aligned reads, similarly showed inconsistent evidence of association with AUROC (Figure 5C): the Pearson correlation was weak and not statistically significant (r = −0.10, p = 0.0608), while the Spearman rank correlation was weak, negative, and nominally significant (ρ = −0.12, p = 0.0144), providing only marginal support for a negative association between local sequence diversity and prediction performance.

**Figure 5.**
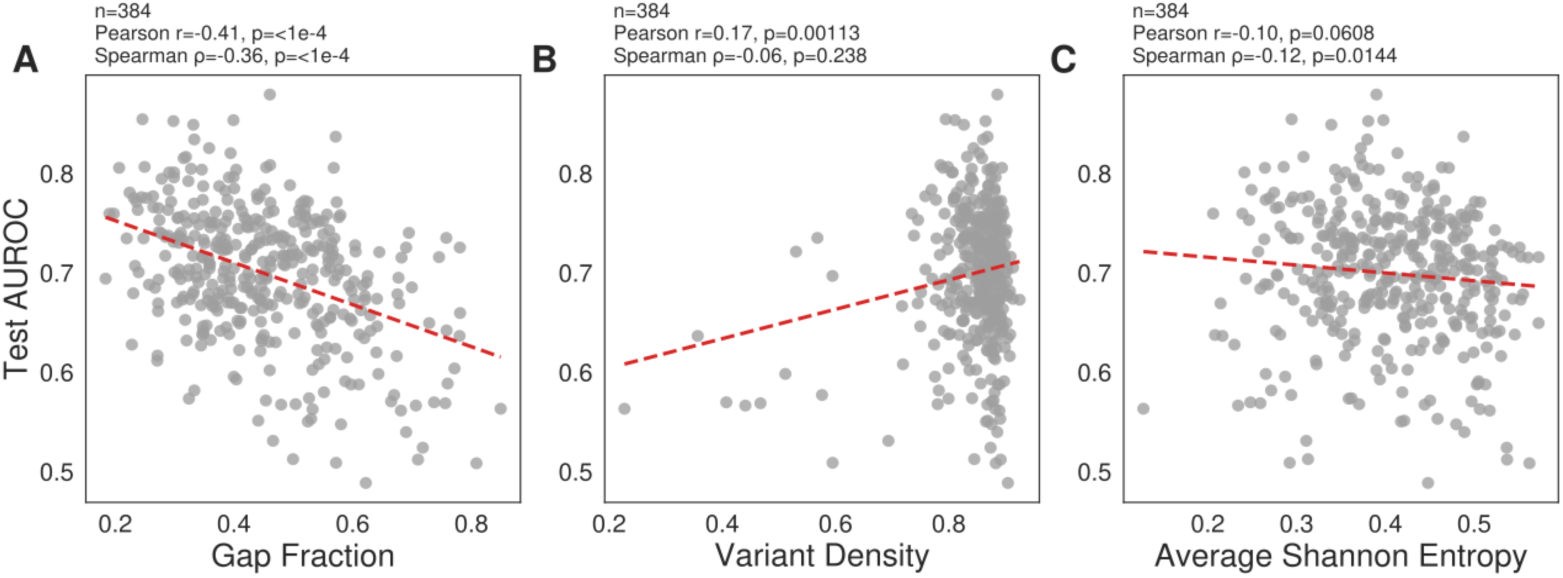
Factors associated with CNN model prediction performance across loci. (A) Relationship between test AUROC and gap fraction across all 384 single-locus CNN models. Higher gap proportions in the input alignment were associated with lower prediction performance (Pearson r = −0.41; Spearman ρ = −0.36; both p < 10⁻⁴), consistent with missing data reducing model accuracy. (B) Relationship between test AUROC and variant density. Variant density showed a weak positive linear association with AUROC (Pearson r = 0.17, p = 0.001) that was not supported by rank correlation (Spearman ρ = −0.06, p = 0.24), suggesting this association is driven by a small number of low-density loci. (C) Relationship between test AUROC and average per-position Shannon entropy. Higher entropy values were weakly associated with lower AUROC (Spearman ρ = −0.12, p = 0.014; Pearson r = −0.10, p = 0.061). Red dashed lines indicate linear regression fits, and inset statistics report Pearson correlation coefficients (r) and Spearman rank correlations (ρ) with associated p-values.

### Effect of read depth on performance

To assess the effect of sequencing depth on model performance, we evaluated a subset of the 10 top-performing CNN locus models across a range of downsampled read depths (from full depth down to 1/32 of full depth; Supplemental Figure 3). Averaged across these 10 loci, mean AUROC decreased from 0.880 at full depth to 0.796 at 1/32 of full depth, and mean accuracy decreased from 0.747 to 0.699 over the same range. An initial decline between full and half depth was followed by a plateau across the intermediate depth fractions (1/2 through 1/16), with the largest single-step decline occurring only at the most extreme reduction tested (1/16 to 1/32 of full depth). Even at this lowest depth, mean performance remained well above the random-classification baseline (AUROC and accuracy of 0.5), indicating that single-locus CNN models remain informative under substantially reduced sequencing depth.

The RF model showed a different pattern across the same depth range (Supplemental Figure 3C). Rather than declining steadily with depth, RF accuracy and AUROC fluctuated within a comparatively narrow band: accuracy ranged from 0.706 to 0.765 and AUROC ranged from 0.749 to 0.870 across all depth fractions tested, with the lowest values occurring at intermediate depths (1/8 for accuracy, 1/16 for AUROC) rather than at the lowest depth evaluated. At 1/32 of full depth, RF accuracy (0.743) and AUROC (0.830) were close to their full-depth values (0.765 and 0.870, respectively), indicating that RF performance is comparatively insensitive to sequencing depth across the range tested. More broadly, these results suggest that a smaller number of high-quality target-capture reads may be sufficient to achieve reasonable predictive performance, particularly when locus-level predictions are aggregated by the RF model.

### Model validation on *Lygodium* dataset

To evaluate performance on a separate empirical dataset, we applied the previously trained models to 60 *Lygodium* samples with ploidy determinations based on chromosome counts (see Methods). Identities and per-model performance of the specific loci and locus pairs evaluated are provided in Supplemental Figure 4.

Across the top-performing models of each type, single-locus CNN models showed the lowest performance (AUROC range: 0.32–0.59, mean = 0.51 ± 0.10; accuracy range: 0.41–0.67, mean = 0.56 ± 0.09). Ensemble models achieved higher AUROC (AUROC range: 0.42–0.74, mean = 0.58 ± 0.14; accuracy range: 0.48–0.58, mean = 0.54 ± 0.04), and hybrid models performed comparably to ensemble models (AUROC range: 0.42–0.74, mean = 0.59 ± 0.14; accuracy range: 0.43–0.59, mean = 0.53 ± 0.07) (Figure 6A, B). However, the newly trained random forest model, which summarizes CNN probabilities across all available loci per sample, outperformed all other approaches (accuracy = 0.782, AUROC = 0.781; 55 labeled samples, 91 samples with usable feature profiles), indicating that aggregating complementary signal across a larger number of loci can further improve cross-taxon generalization beyond pairwise ensembling alone. For comparison, nQuire achieved an accuracy of only 0.281 (full) and 0.263 (denoised) on the *Lygodium* dataset, performing well below chance, while nQuack achieved an accuracy of 0.643, below both the random forest and the best individual CNN models. The relatively strong performance of nQuack was greatly influenced by the high proportion of diploids in the *Lygodium* dataset as it again appears to have difficulty identifying polyploids.

**Figure 6.**
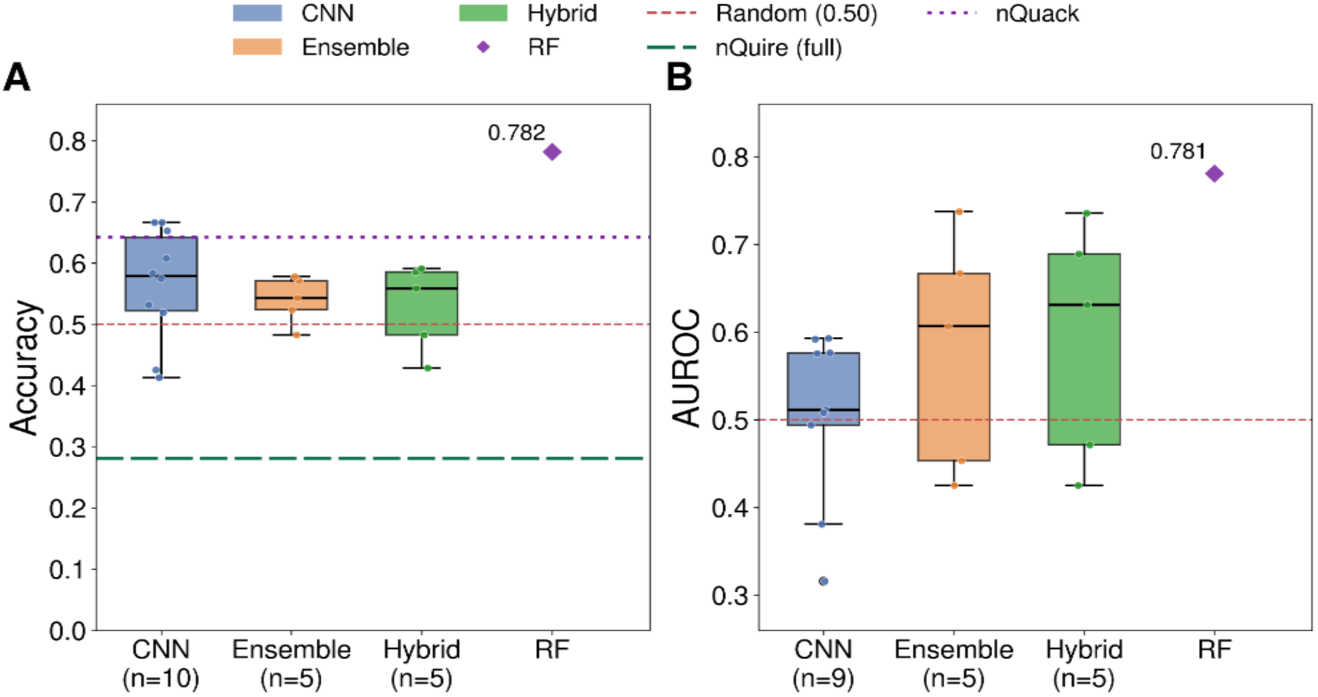
Generalization of trained models to the independent *Lygodium* dataset. (A) Classification accuracy and (B) AUROC of the top-performing single-locus CNN (n = 10 for accuracy, n = 9 for AUROC), ensemble (n = 5), and hybrid (n = 5) models applied to 60 *Lygodium* samples with known ploidy; because locus recovery varied among samples, not every model could be evaluated on all 60 samples. CNN models are shown in blue, ensemble models in orange, and hybrid models in green; points show individual models. The purple diamond indicates the random forest (RF) model, which aggregates locus-level CNN probabilities across all available loci per sample (accuracy = 0.782, AUROC = 0.781). The red dashed line indicates random classification performance (0.50); in (A), the green dashed and purple dotted lines indicate the accuracy of nQuire applied to full (undenoised) read data (0.281) and nQuack (0.643), respectively. One CNN model (L138) could be evaluated on only three samples and lacks an AUROC because all three were diploid. Ensemble and hybrid models generally achieved higher AUROC than single-locus CNN models, and the RF model outperformed all other approaches, suggesting that aggregating predictions across many loci improves robustness.

To better understand the differences in classification performance among model types, we examined their predicted-probability distributions relative to the 0.5 decision threshold (Supplemental Figure 5; n = 53 CNN, n = 48 ensemble, and n = 47 hybrid per-sample predictions). For each sample, we averaged the probabilities from the top-performing CNN models (up to 10) and from the top-performing ensemble pairs (up to 5), and computed a hybrid value as the average of these two per-sample means. The CNN predictions were concentrated below 0.5 (i.e., toward diploid), with only 3.8% of per-sample mean probabilities exceeding 0.5, despite an observed polyploid prevalence of 34.0%. This suggests that the single-locus CNN models tended to classify *Lygodium* samples as diploid, potentially reflecting poor probability calibration on this dataset. In contrast, ensemble predictions were more dispersed and frequently exceeded the classification threshold, with 56.2% of per-sample mean probabilities above 0.5 against an observed prevalence of 31.2%. The hybrid values produced the most balanced threshold behavior, with 29.8% exceeding 0.5, closely matching the observed polyploid prevalence of 31.9%. These results suggest that combining CNN and ensemble predictions may partially offset their opposing biases, bringing the proportion of polyploid calls closer to the observed prevalence.

### Autopolyploid vs allopolyploid

Finally, we examined whether our trained models exhibit distinct prediction performance when applied to autopolyploid versus allopolyploid samples, by subsetting the *Lygodium* dataset into autotetraploid vs. diploid and allotetraploid vs. diploid comparisons, based on the classifications made by Pelosi et al. (2024) using phylogenomic approaches. In both comparisons, diploid samples substantially outnumbered the tetraploid subset, which would artificially inflate the accuracy of a model that simply predicted "diploid" for every sample, independent of any genuine discriminative ability. For example, a naive classifier that always predicted "diploid" — regardless of model output — would achieve 79.5–80.5% accuracy on the autotetraploid vs. diploid comparison and 85.4–86.8% accuracy on the allotetraploid vs. diploid comparison. These ranges reflect the slightly different sample sizes available to the CNN, ensemble, hybrid, and RF models due to differing per-sample locus coverage, not a difference in model performance. We report bootstrap accuracy (see Methods) for each model type in Supplemental Figure 6A, B for completeness. Notably, observed accuracy fell below this naive baseline for all deep-learning model types in both comparisons (CNN: 0.751 and 0.805; ensemble: 0.465 and 0.554; hybrid: 0.652 and 0.703, for autotetraploid and allotetraploid discrimination, respectively), confirming that these naive baselines are already high without reflecting any real discrimination. The RF model achieved accuracy of 0.779 and 0.857 for autotetraploid and allotetraploid discrimination, respectively — approaching the naive baselines but still reflecting genuine discrimination as confirmed by AUROC. We therefore treat AUROC — which is threshold-independent and not biased by class imbalance — as the primary metric for this comparison (Supplemental Figure 6C, D).

Bootstrap-aggregated CNN AUROC was higher for autotetraploid vs. diploid discrimination (0.657; 95% CI: 0.47–0.83) than for allotetraploid vs. diploid discrimination (0.372; 95% CI: 0.13–0.63); the point estimate for allotetraploid discrimination fell below the random-classification value of 0.5, suggesting that single-locus CNN models do not reliably distinguish allotetraploids from diploids. Ensemble models showed the opposite pattern, with higher AUROC for allotetraploids (0.722; 95% CI: 0.51–0.92) than for autotetraploids (0.361; 95% CI: 0.11–0.63). The hybrid approach followed the same direction as the ensemble models, with higher AUROC for allotetraploids (0.675; 95% CI: 0.45–0.87) than for autotetraploids (0.400; 95% CI: 0.15–0.66). In contrast, the RF model achieved the highest AUROC of any method for allotetraploid discrimination (0.856; 95% CI: 0.69–0.98) and matched CNN performance for autotetraploid discrimination (0.671; 95% CI: 0.47–0.85), making it the most consistent model type across both subgroups. Because these comparisons include only 10 autotetraploid and 6 allotetraploid samples, the confidence intervals are wide and overlap between subgroups, and these differences should be interpreted with caution.

These contrasting patterns raise the possibility that CNN and ensemble models capture different aspects of the ploidy signal: individual CNN models, which rely on single-locus read patterns, may be more sensitive to the dosage effects characteristic of autopolyploidy, while ensemble models, which integrate information across locus pairs, may be better suited to detecting the inter-locus divergence patterns associated with allopolyploidy. The RF model, which aggregates CNN predictions across all available loci, may combine both signals. Testing these hypotheses will require larger samples of autopolyploids and allopolyploids with known origins.

## DISCUSSION

### Performance of sorAI

The sorAI machine learning modelling approaches infer polyploidy from GoFlag targeted capture data of ferns with a degree of accuracy that surpasses that of more general statistical approaches like nQuire (Weiß et al., 2018) or nQuack (Gaynor et al., 2024). Perhaps more importantly, rather than simply being a black box to infer ploidy from complex, previously undetected patterns, we demonstrate how ML approaches can be used to provide insights into the data features most important for determining ploidy and the potential shortcomings of other approaches to infer ploidy from HTS data.

There appear to be some broad associations with ploidy level across all the loci. For example, polyploids tend to have higher overall heterozygosity across all loci than diploids (Supplemental Figure 7). However, these associations are not clear or distinct enough to make them reliably diagnostic about ploidy level. While the ML models leverage targeted capture and chromosome count data from hundreds of fern specimens to build the training set, the CNN models demonstrate that it is possible to infer ploidy with high accuracy from even a single locus (Figure 2A). This is reminiscent of early molecular approaches that determined ploidy in ferns using allozymes or few loci (e.g., Haufler, 1985). Yet there is high variance in the performance of CNN models among loci; in fact, models developed from many loci are no better, and occasionally worse, than random guesses (Figure 2). This suggests that focusing on the most informative loci may provide more clarity than summarizing patterns from all loci. Our ML models provide such an approach to highlight these most informative loci.

Although the single-locus CNN models often perform well, in some cases, their performance may falter. For example, CNN models appear to have decreased accuracy when sequence coverage or data quality is lower, as seen in the experiments decreasing the sequencing depth (Supplemental Figure 3) or in the analyses of the *Lygodium* dataset, in which there was below average recovery of loci from GoFlag probe set (Figure 6). The performance of the ensemble models and the Random Forest model show that adding information from multiple loci may improve upon and provide more robust estimates of ploidy than the single locus CNN models. In most cases we examined (more than 92% of locus pairs), the ploidy estimates from the two-locus ensemble models improve upon the estimates of the single-locus CNN models using their constituent loci (Figure 3). The RF models maintain an accuracy of approximately 71–78% on the original test set (76.5%), in the difficult Lygodium dataset (78.2%; Figure 6), and across all sequencing depths tested, including 1/32 of the original depth (74.3%; Supplemental Figure 3). Thus, our experiments show that the information about ploidy is unevenly distributed among loci, but leveraging information from multiple loci provides the most stable estimates of ploidy.

### What information is sorAI using

With any ML approach, a natural question is what features of the data most influence the results? Perhaps not surprisingly, the amount of missing data appears to be the greatest predictor of performance of a locus in the CNN models (Figure 5A). There also is a weak negative correlation of the Shannon entropy of the read alignment for a locus and the resulting AUROC score (Figure 5C), suggesting that more complex alignments are not as informative for predicting ploidy. However, it remains difficult to determine a priori which loci will perform best in the CNNs. Although it is intuitive that combining information from multiple loci in the ensemble models improves upon predictions from the single-locus CNN models (Figure 3), the performance of the ensemble models is difficult to predict from the CNN models. For example, combining the loci with the best CNN models does not generate the best ensemble model. This also suggests that the CNN models and the ensemble models may use different features of the loci. The difficulty of predicting the performance of different loci emphasizes the potential use of an approach that can evaluate the performance of many loci using a large training set.

Furthermore, our randomization experiments (i.e., where rows and columns are shuffled in read alignments) demonstrate that the ML models recognize and capitalize on variation among alignment sites to accurately infer ploidy. Randomly swapping alignment columns greatly reduces the accuracy of the ML approach (Figure 4B), indicating that some sites in the training data contain more information about ploidy than others. This is in contrast to the approach of nQuire and nQuack, in which all alignment sites equally contribute to the site frequency distributions, and information about their order is discarded. Furthermore, by randomly shuffling the nucleotides from different reads from the same sample, we find that maintaining the covariance among sites also contributes information about the ploidy of the sample, especially in the ensemble models (Figure 4B). The potential information in the neighboring sites in determining ploidy can be demonstrated in an example with four sequence reads that are each two base pairs in length: “AG”, “AT”, “CG”, and “CT”. A method based on read frequencies that treats each site independently, like nQuire or nQuack, would interpret these reads coming from a diploid because of the frequency of each allele in the two sites is 50% (i.e., 50% A and 50% C in site 1; 50% G and 50% T in site 2). However, if the sites are considered together, accounting for the covariance among sites, then these sequences would be interpreted as four distinct alleles, suggesting they did not come from a diploid. By using information about each site and accounting for the states of the neighboring sites in each read, the ML approach achieves its greatest accuracy (Figure 4B).

### Limitations of Machine Learning Models

While the ML approach overall performed well, our experiments suggest we still might expect error rates of 22–29% using the RF model. So why aren’t we able to achieve higher accuracy? One potential source of error is the training set. To assemble the training set, we first determined whether a sample was from diploid or polyploid based on species chromosome counts; however, these can be misleading. For example, some fern taxa with high chromosome numbers are actually diploid (e.g., Haufler and Soltis, 1986; Wolf et al., 1987; Rizzieri et al., 2026). Also, we do not have chromosome counts from the specific fern samples from which we have targeted capture data. There are numerous cases of fern species complexes that include both diploid and polyploid individuals (e.g., Manton, 1950; Wagner, 1954; Hanušová et al., 2019; Pelosi et al., 2024), and therefore, it is possible that the sequence sample came from an individual with a different ploidy than the individual from which the chromosome count was obtained.

An alternate approach to assemble large and accurate training sets would be to generate them through simulations. This is conceptually similar to the approach of Sutherland et al. (2024) to infer ancient whole genome duplications from Ks plots using ML models trained on simulated datasets. Indeed, there are new tools that can simulate complex polyploid scenarios (e.g., Blischak et al., 2023; Conant, 2023; Dunn and Sethuraman, 2024, 2025). By simulating specific polyploid evolution scenarios, it also might be possible to extend the current model to distinguish among different ploidy levels or distinguish allopolyploids from autopolyploids. While such an approach is promising, even our best models of sequence evolution likely fail to incorporate realistic complexity (Trost et al., 2024), and such training sets may have their own biases.

There also likely are inherent limitations to the information about ploidy in target capture datasets that may limit any approach to infer ploidy. The GoFlag probe set was designed from alignments of single copy nuclear genes identified by the 1KP transcriptome sequencing project (One Thousand Plant Transcriptomes Initiative, 2019). Focusing on single copy loci often is desirable for phylogenetic analyses because they presumably minimize errors caused by undetected paralogy. However, the lack of duplicate copies in these genes likely reflects strong selection to eliminate extra copies after a gene duplication or polyploidy event, erasing the evidence of polyploidy. Although some studies have focused on the use of nQuire for inferring ploidy using target capture data (Viruel et al., 2019, 2023), originally both nQuire and nQuack were described for use more broadly with genomic data, which likely would include more multi-copy genes. Our ML approach can easily be adapted to any short read HTS datasets, but since targeted capture is so widely adopted for phylogenetics, there is value in exploring its potential for inferring ploidy. One future approach might use the growing abundance of genomic and transcriptomic data to determine the genes that are most indicative of ploidy level and add probes that cover these genes in future targeted capture probe sets. For example, in the ferns Pelosi et al. (2022) identified convergence in the patterns of genes retained in duplicate across independent paleopolyploidy events; confirming the duplicate nature of these loci in newly generated chromosome-scale genomes (e.g., Li et al., 2018; Rizzieri et al., 2026; Xia et al., 2026) could identify these ploidy-informative loci.

The *Lygodium* dataset provided a challenge for all methods to infer ploidy, and also points the direction towards possible approaches to improve performance in the future. The generally lower performance using nQuack, nQuire, and the CNN models with *Lygodium* (Figure 6) likely are due, at least in part, to the higher levels of missing data than occur in many targeted capture datasets (see Pelosi et al., 2024, on average only 330 biallelic sites were retained after filtering with nQuire). *Lygodium* appears to contain a combination of autopolyploids and allopolyploids (Pelosi et al., 2024), and this also may contribute to reduced accuracy. Our ML training set does not distinguish between autopolyploids and allopolyploids. However, if there are relatively few autopolyploid ferns, as suggested in Wagner and Wagner (1980; but see Barker et al., 2016), the ML models may have little training on patterns associated with autopolyploidy. While the accuracy of the CNN, Ensemble, and Hybrid approaches in *Lygodium* is lower than we see in the test set across ferns (Figure 6), the relatively higher AUROC scores indicate that the relative probability of ploidy output from the models is still generally higher in polyploids than in diploids. Thus, in theory, we could likely improve upon the accuracy by using a different probability cutoff for inferring a polyploid. However, it is difficult to determine the optimal cutoff a priori. The RF approach provides a more workable solution that maximizes the performance of the ML approaches by combining insights from all CNN models. In fact, it appears to perform as well as on the original test set on both allo-and auto-polyploids, with little evident bias. More investigation is needed to broadly determine the best approaches to infer ploidy from the ML results across various datasets and how to best quantify uncertainty associated with ML approaches. No matter the method one is using to infer ploidy, the *Lygodium* example points towards the importance of uncertainty (Beaulieu and O’Meara, 2025).

Unlike nQuire or nQuack, the specific models we present in this paper were trained on a specific probe set in one large land plant clade and may not be easily applied to other clades or probe sets. Although the GoFlag 408 probe set was designed to cover all flagellate land plants, including all gymnosperms (Breinholt et al., 2021), each clade poses its own challenges. For example, the bryophytes likely will have very different patterns of sequence variation associated with polyploidy than ferns simply because most DNA extractions used gametophyte rather than sporophyte tissue. Polyploidy is rare in gymnosperms; *Ephedra* includes by far the most polyploid taxa (see Ickert-Bond et al., 2020), yet, *Ephedra* samples have extremely low locus recovery rates with GoFlag sequencing, likely due to probe design (Ickert-Bond and Burleigh, unpublished). Beyond these more technical issues, ultimately our ability to generalize these models to different clades depends on the degree to which the same loci or same sites associated with ploidy in ferns also would be associated with ploidy in other clades. Notably, the rediploidization process in ferns appears to differ from that in other clades (e.g., angiosperms; Haufler, 1987; Kinosian, 2026); the biological signal of polyploidy that the ML approach implemented here is likely not the same in other lineages. It is straightforward to train new ML models for inferring ploidy from alignments of sequence reads using our approach in different clades and probe sets. This investment is more computationally intensive than using more easily generalizable approaches such as nQuire or nQuack; however, they offer the advantage of being tailored to potentially heterogeneous or idiosyncratic patterns associated with ploidy that may occur in different clades or loci. Once the models are developed, they are quick and simple to run, even on a laptop.

### Best practices for applying this method to new datasets

Based on our results, we outline several practical recommendations for researchers applying this approach to new targeted capture datasets. First, prediction accuracy for the CNN or ensemble models depends on input data quality: our downsampling analysis (Supplemental Figure 3) and the relationship between AUROC and alignment gap fraction (Figure 5A) both indicate that higher-depth, higher-quality target-capture data — comparable to most GoFlag loci used here — yield more reliable predictions with the CNN or ensemble models, while lower depth or gap-rich alignments degrade model performance. However, the random forest (RF) model appears to be relatively robust to data quality, achieving approximately 71–78% accuracy across all sequencing depths tested (74.3% at 1/32 of full depth) and 78.2% in the Lygodium dataset, where the accuracy of the CNN and ensemble models is reduced. Although it is possible to achieve potentially higher levels of accuracy in high quality datasets from single locus CNN models or two locus ensemble models, unless the user is confident of the quality and abundance of their data, in most cases we recommend using the predictions of the RF model.

We did not test the models on samples from taxa outside of the training set (i.e., non-ferns) although many or all these loci can be amplified in other groups with existing probe sets. The GoFlag408 flagellate land plant probe set can be used to generate targeted capture data from any non-angiosperm land plants (Breinholt et al., 2021), and there exist GoFlag probe sets to amplify the same loci in angiosperms (Endara et al., in prep.) and algae (Sigel and Burleigh, in prep.). There also is much overlap in the loci from the Angiosperm 353 probe set (Johnson et al., 2019) and the GoFlag probe sets (Breinholt et al., 2021; Endara et al., in prep.). In those cases, we suspect that individual locus-level predictions will be substantially noisier, and we recommend against relying on any single locus (i.e., using CNN or ensemble models). Where feasible, users with their own ploidy verified reference samples should re-train locus-specific and ensemble models on their own data. When retraining is not feasible, aggregating information across many loci — particularly using the random forest combiner, may provide the most robust predictions for taxa outside the training distribution.

Finally, we recommend that predicted ploidy be treated as a probabilistic rather than categorical outcome in downstream applications, or minimally that downstream analyses account for potential error in the ploidy designations output from these models. Because our models output a continuous probability of polyploidy rather than a hard classification, this probability itself can serve as a built-in measure of prediction confidence, and downstream analyses (e.g., comparative phylogenetic studies that incorporate inferred ploidy) can propagate this uncertainty rather than treating each call as certain. Additional uncertainty estimates could be obtained by subsampling reads to generate a bootstrap distribution of predictions per sample, or through other established machine-learning approaches for uncertainty quantification. We did not implement these here, but see them as a valuable direction for future work.

## Acknowledgments

This work was funded in part by U.S. National Science Foundation (GoFlag; DEB-1541506). J.A.P. was supported by USDA NIFA Postdoctoral Fellowship #2024-67012-43394. F.R. and J.Z. were supported by NIH grant R35GM15490.

## Author Contributions

All authors designed the experiments, performed the analyses, and wrote the paper.

## Data Availability Statement

The sorAI code, including the preprocessing, training, and inference pipelines and the trained random forest model, is available on GitHub at https://github.com/Fahimehgit/sorai. Trained model weights are available on Hugging Face at https://huggingface.co/Fahimehr140/sorai. Per-sample model predictions and performance metrics are provided in Supplemental Table S1.

## Supplemental Figures

**Supplemental Figure 1.**
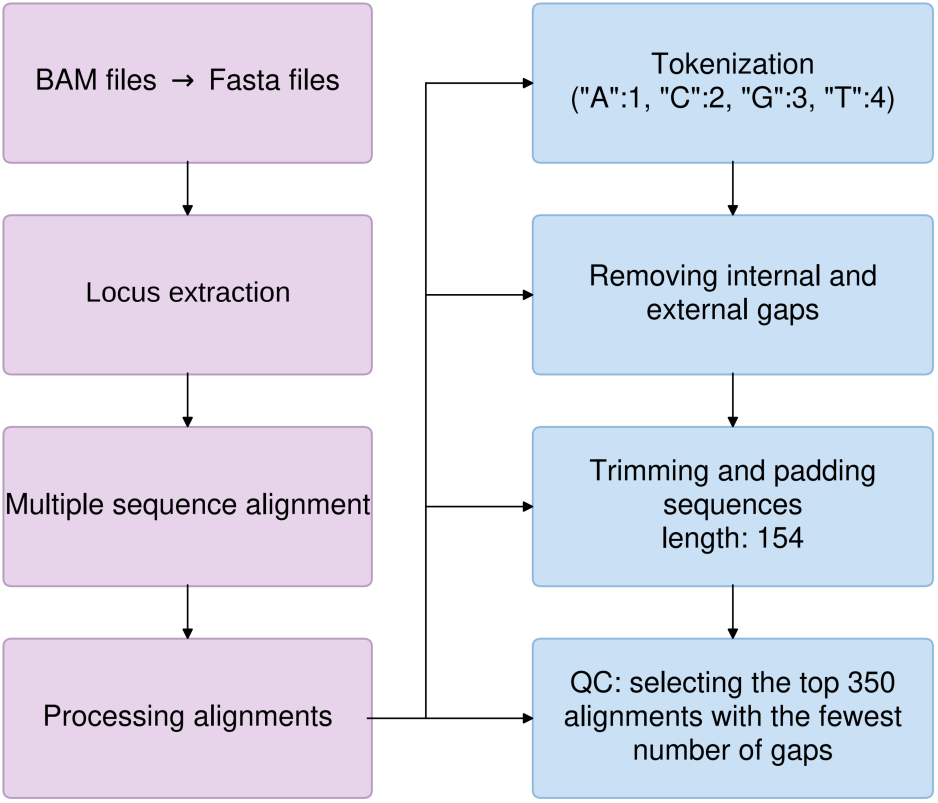
Detailed bioinformatics pipeline for processing target-capture reads. For each sample and locus, BAM files were converted to FASTA format, followed by locus-specific extraction, multiple sequence alignment, and alignment processing. Alignment processing included tokenization of nucleotides (A = 1, C = 2, G = 3, T = 4, gap/other = 0), removal of internal and external (leading/trailing) alignment gaps, trimming and padding of sequences to a fixed length of 154 positions, and a final quality-control step retaining the 350 reads per locus with the fewest gap characters (padding with the gap character when fewer than 350 reads were available), consistent with the pipeline summarized in Figure 1C and described in Methods.

**Supplemental Figure 2.**
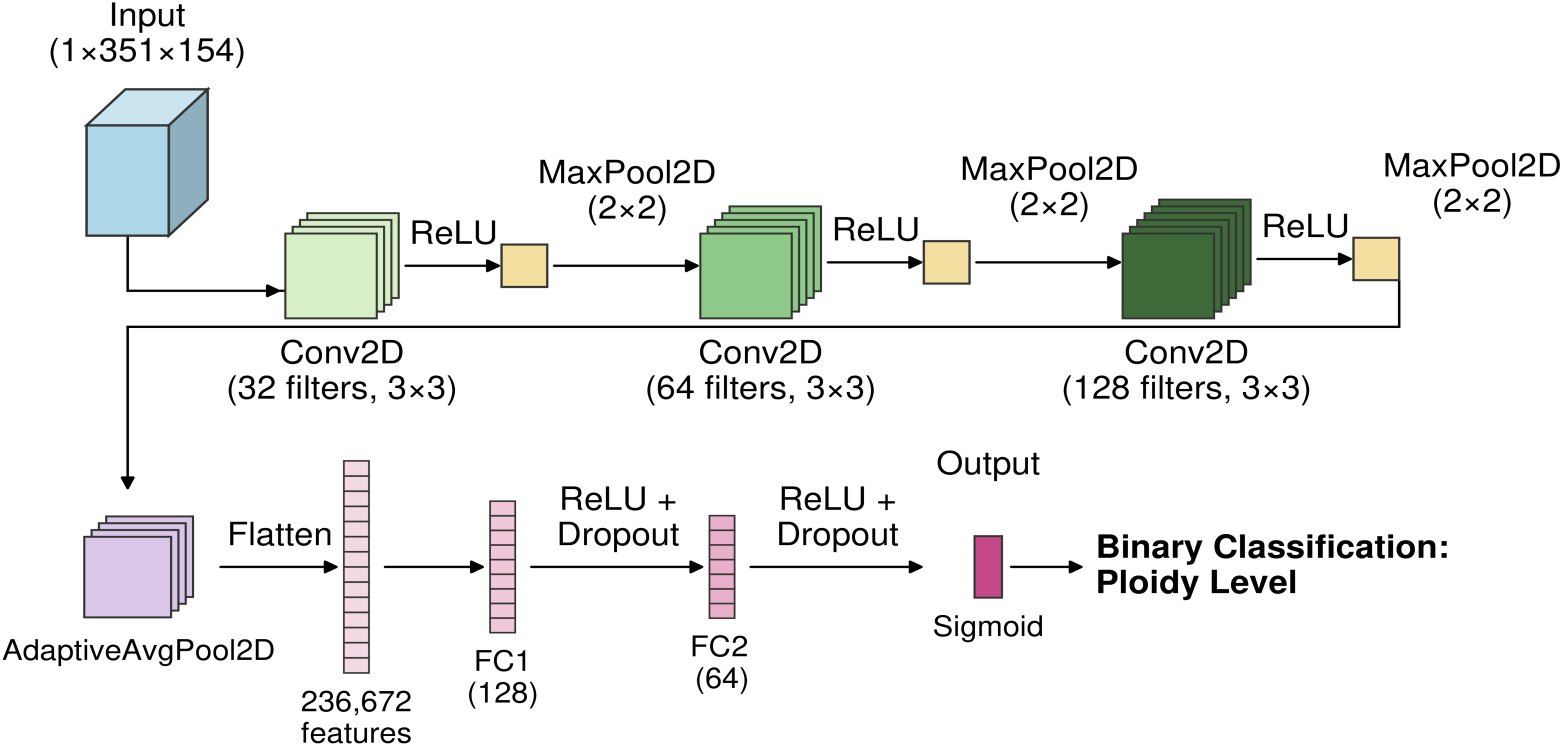
Representative CNN architecture for ploidy classification. Schematic illustrating the general design of the single-locus CNN models. The model input consists of a single-channel array representing aligned reads by alignment position (1×351×154, corresponding to 1 reference sequence plus 350 reads). In this example, the input is passed through three convolutional blocks (32, 64, and 128 filters, respectively; 3×3 kernels), each followed by ReLU activation and 2×2 max pooling. The resulting feature maps are reduced via adaptive average pooling and flattened (236,672 features), then passed through two fully connected layers (FC1, 128 units; FC2, 64 units), each followed by ReLU activation and dropout. A final sigmoid output layer produces the predicted probability of polyploidy. Layer sizes are illustrative: the number of convolutional layers (1–3), filter counts (16–32 per layer), kernel size (3×3 or 4×4), activation function, pooling type, and fully connected layer width (64–128 units) were optimized independently for each locus via Optuna (Supplemental Table S1), so the 64-and 128-filter blocks shown here fall outside the searched range and do not correspond to any single trained model.

**Supplemental Figure 3.**
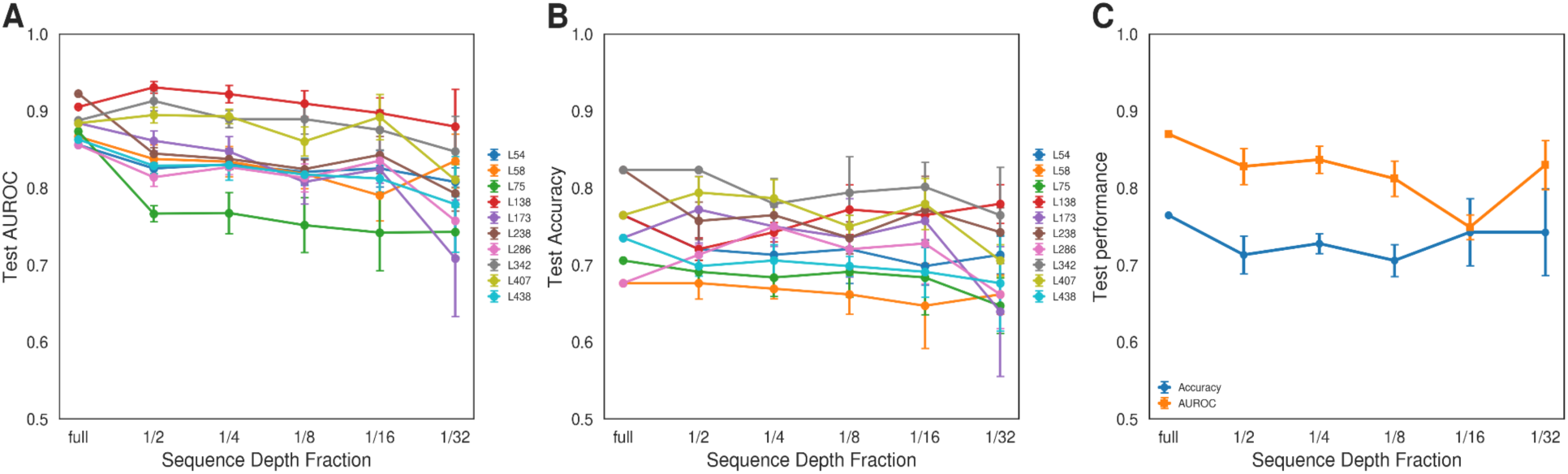
Effect of sequencing depth on CNN and RF model performance. (A) Test AUROC and (B) test accuracy across sequencing-depth fractions (full, 1/2, 1/4, 1/8, 1/16, and 1/32) for the ten top-performing single-locus CNN models used for depth-sensitivity benchmarking. For each locus and depth fraction, non-reference reads were randomly subsampled to the target depth and randomly reordered to minimize order-dependent bias; predictions were repeated across four replicate runs, and points show mean performance with error bars indicating ±1 SD across replicates. (C) Random Forest (RF) performance (accuracy and AUROC) across the same depth fractions, using the same random forest model and the same replicate-based depth-reduction and read-reordering procedure.

**Supplemental Figure 4.**
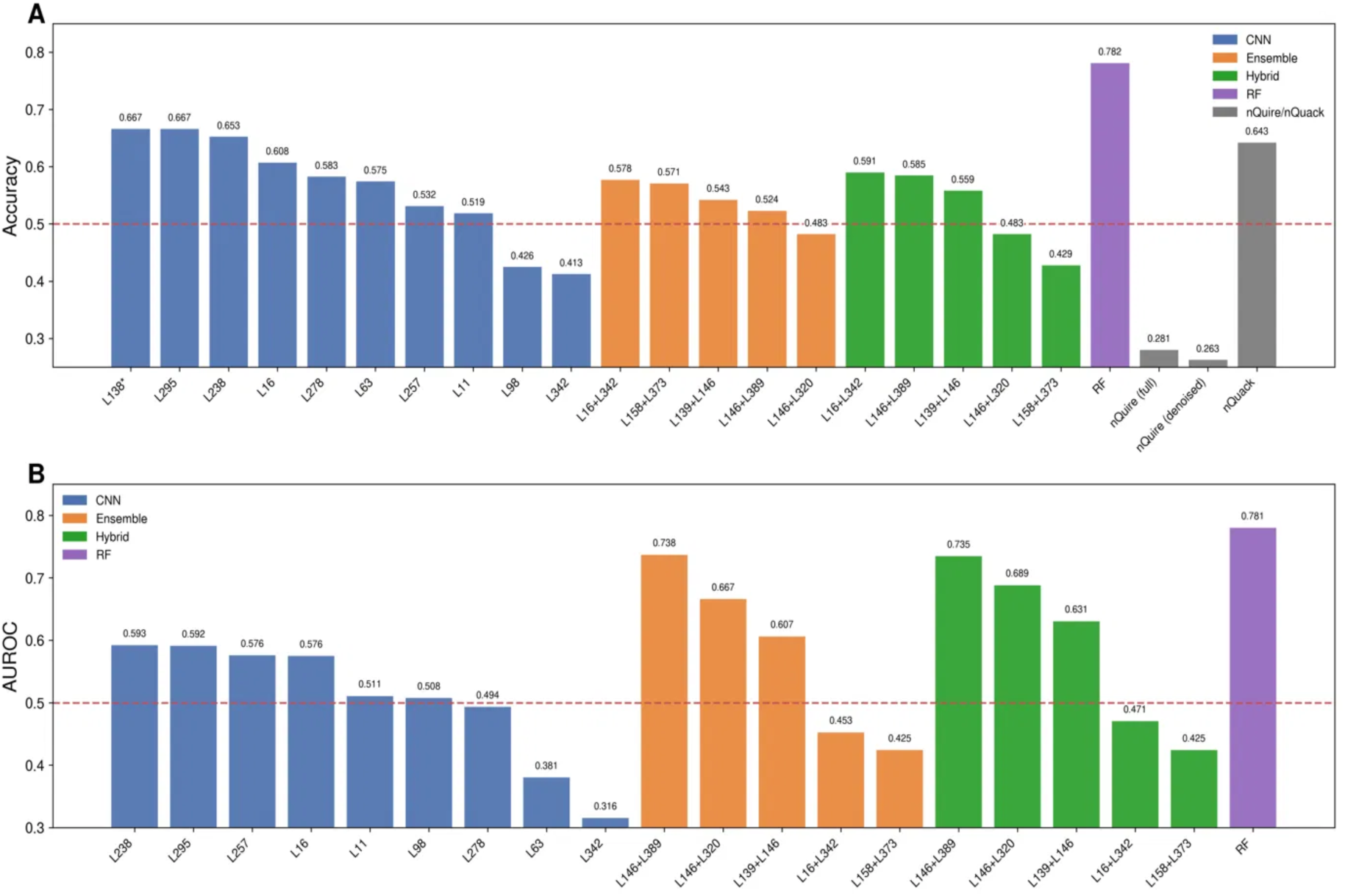
Performance of individual CNN, ensemble, hybrid, and random forest models on the *Lygodium* dataset. (A) Accuracy and (B) AUROC for the top-performing models applied to 60 *Lygodium* samples with known ploidy, grouped by model type: single-locus CNN models (blue), ensemble models combining pairs of loci (orange), hybrid models averaging CNN and ensemble probabilities (green), the random forest model aggregating locus-level CNN probabilities across all available loci (purple), and the allele-frequency–based methods nQuire and nQuack (gray; accuracy only, as these methods return ploidy calls rather than probabilities). The random forest model achieved the highest accuracy (0.782) and AUROC (0.781). The best-performing single-locus CNN model reached an accuracy of 0.667 (L295, evaluated on 42 samples). *L138 reached the same accuracy but could be evaluated on only three samples, all diploid, so its AUROC is undefined, and it is omitted from (B). nQuack achieved an accuracy of 0.643, while nQuire performed below chance (full: 0.281; denoised: 0.263). Red dashed lines indicate random classification performance (0.50).

**Supplemental Figure 5.**
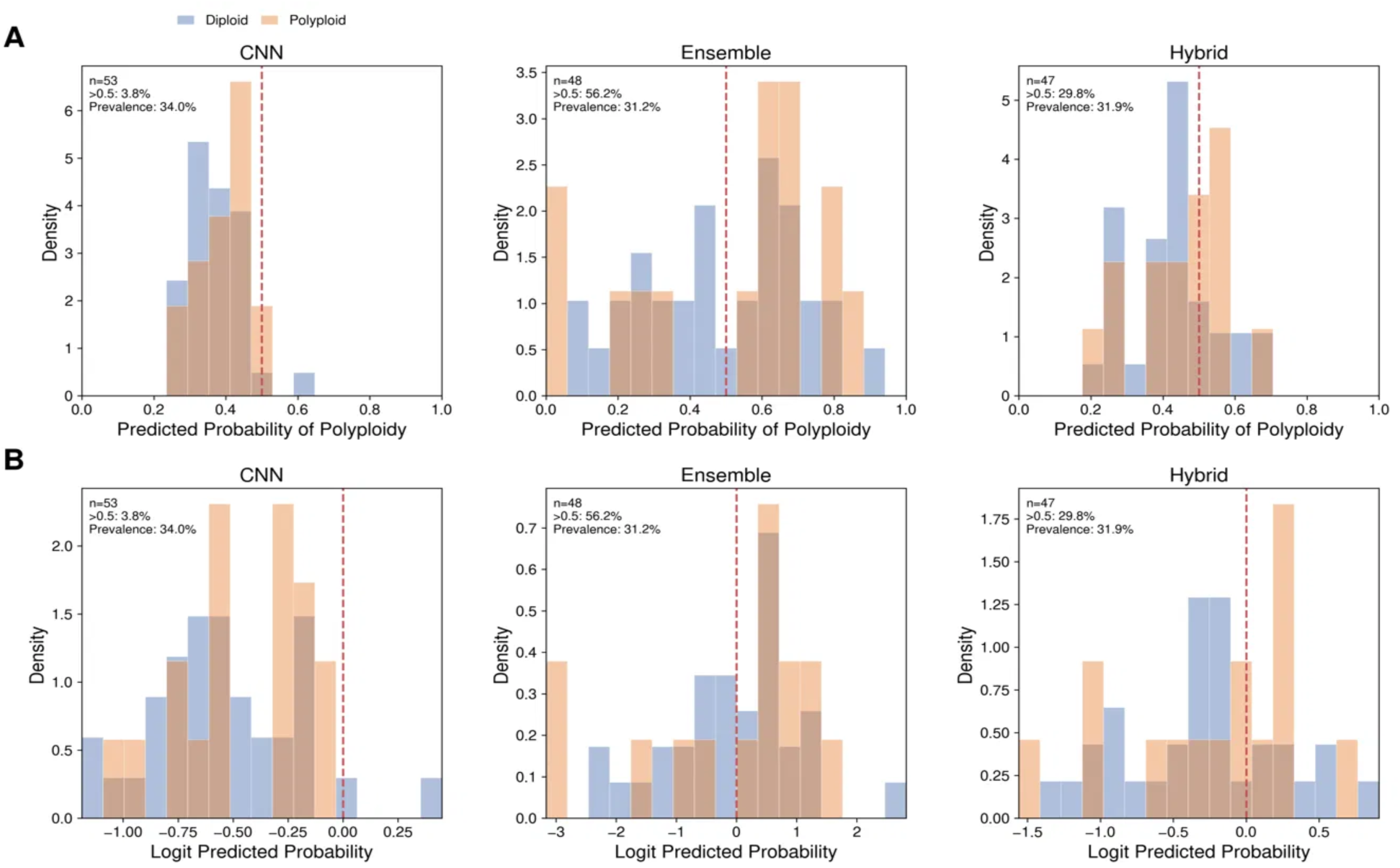
Distributions of predicted probabilities on the *Lygodium* dataset. (A) Distribution of per-sample predicted probabilities of polyploidy for the CNN (n = 53), ensemble (n = 48), and hybrid (n = 47) approaches, colored by true ploidy status (diploid = blue, polyploid = orange). For each sample, the CNN value is the mean probability across the top-performing CNN models that could score that sample (up to 10), the ensemble value is the mean across the available top-performing ensemble pairs (up to 5), and the hybrid value is the average of the sample’s CNN and ensemble values. Histograms are normalized within each ploidy class, and the dashed vertical line indicates the 0.5 classification threshold. (B) The same values on the logit scale, with the dashed line indicating logit(p) = 0. CNN predictions were skewed toward diploid calls, with only 3.8% of samples exceeding the 0.5 threshold despite an observed polyploid prevalence of 34.0%. Ensemble predictions exceeded the threshold more often than the observed prevalence (56.2% vs. 31.2%), whereas the proportion of hybrid predictions above the threshold most closely matched the observed prevalence (29.8% vs. 31.9%).

**Supplemental Figure 6.**
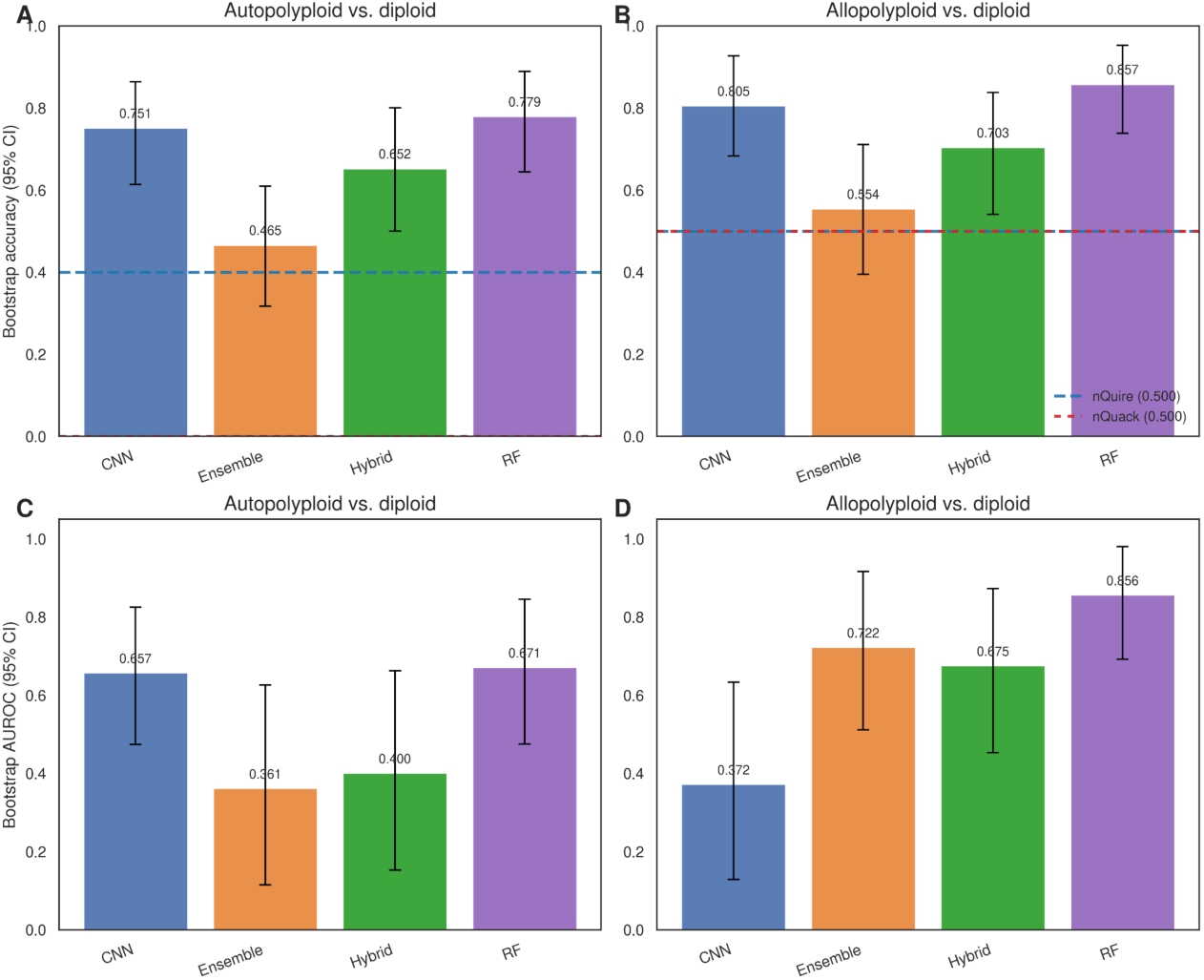
Model performance on *Lygodium* autopolyploid and allopolyploid samples relative to diploids. (A, B) Bootstrap accuracy (mean ± 95% CI) for CNN, ensemble, hybrid, and RF models (per-sample values computed as in Supplemental Figure 5) evaluated on (A) autopolyploid vs. diploid and (B) allopolyploid vs. diploid samples. Among the CNN, ensemble, and hybrid approaches, CNN achieved the highest accuracy (autopolyploid: 0.751; allopolyploid: 0.805), while RF was comparable or better (autopolyploid: 0.779; allopolyploid: 0.857). Ensemble models performed worst (0.465 and 0.554, respectively), and hybrid models were intermediate (0.652 and 0.703, respectively). (C, D) Bootstrap AUROC (mean ± 95% CI) for the same models and comparisons. CNN AUROC was substantially higher for autopolyploid vs. diploid discrimination (0.657) than for allopolyploid vs. diploid discrimination (0.372), while ensemble and hybrid models showed the opposite pattern, performing better on allopolyploid discrimination (ensemble: 0.722; hybrid: 0.675) than on autopolyploid discrimination (ensemble: 0.361; hybrid: 0.400). RF achieved the highest AUROC for allopolyploid discrimination (0.856) and matched CNN on autopolyploid discrimination (0.671 vs. 0.657). Dashed lines in (A, B) indicate nQuire and nQuack baselines (autopolyploid: nQuire = 0.4, nQuack = 0.0; allopolyploid: nQuire = nQuack = 0.5).

**Supplemental Figure 7.**
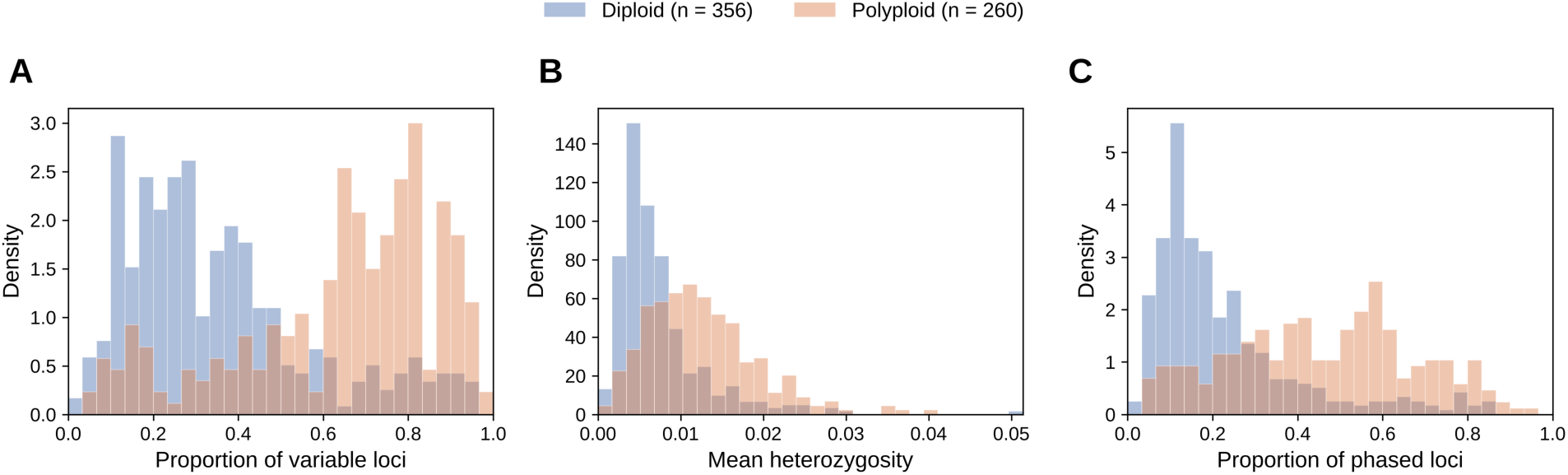
PATÉ phasing summaries for diploid and polyploid samples. Distributions of (A) the proportion of variable loci, (B) mean heterozygosity, and (C) the proportion of phased loci per sample, estimated with PATÉ assuming all samples were diploid, for all 616 samples with complete phasing results (356 diploid and 260 polyploid). Histograms are normalized within each ploidy class. Polyploid samples had a higher proportion of variable loci (median 0.69 vs. 0.29), higher mean heterozygosity (0.0115 vs. 0.0058), and a higher proportion of phased loci (0.47 vs. 0.17) than diploid samples (two-sided Mann–Whitney U tests, all p < 10⁻²⁹). The distributions overlapped, and each statistic used alone as a classifier achieved an AUROC of 0.77–0.80.

